# Characterization of time-dependent and history-dependent mechanical behaviour of human masseter muscle

**DOI:** 10.64898/2026.06.11.731620

**Authors:** Eyad Awad, Noemie Briot, Gregory Chagnon, Raymond Challita, Lucille De Bengy-Puyvallee, Djordje Peric, Mokarram Hossain

## Abstract

The human masseter muscle is one of the primary muscles responsible for mastication and mandibular movement; however, its intrinsic mechanical properties remain insufficiently characterized. In this experimental study, the nonlinear, viscoelastic, and history-dependent behaviour of the human masseter muscle was investigated using ex vivo uniaxial cyclic tensile tests. The masseter muscle samples prepared from fresh and formalin-preserved cadavers were tested under two loading protocols: a continuous stretch protocol with increasing stretch levels and a constant stretch protocol with repeated loading to a fixed maximum stretch. Tests were conducted at two strain rates, and their influences on the mechanical behaviour of the tissue were examined. The effect of formalin preservation was also investigated. The results showed that the stiffness of the tissue increases for formalin-preserved samples. Under cyclic loading, the features including energy dissipation, stress-softening, residual deformation, and cyclic conditioning progressively changed during the initial loading cycles and reached stabilization during the final cycle. These findings provide experimental evidence that the human masseter muscle exhibits nonlinear, viscoelastic, and history-dependent mechanical behaviour under cyclic tensile loading. The experimental data obtained in this study may be used for biomechanical modelling of the human masticatory system and the development of constitutive models for cranio-maxillofacial surgical simulation, prosthetic design, and facial soft-tissue biomechanics.

**Statement of significance:** The masseter muscle is one of the primary muscles of mastication. To address the current gap in craniofacial biomechanics that has largely focused on the mechanical characterization of the masseter muscle based on imaging techniques or monotonic loading, this study quantifies the nonlinear and viscoelastic mechanical response of masseter tissue under cyclic continuous and constant stretch loading, including strain-rate and preservation effects. The results show that the mechanical behaviour of the masseter muscle, including stiffness, hysteresis, stress-softening, and residual strain behaviour, is strongly influenced by strain-rate and formalin preservation. The experimental results provide mechanical data for constitutive modelling of the masticatory system with applications in cranio-maxillofacial surgical simulation, prosthetic design, and facial soft tissue modelling.

## 1. Introduction

The masseter muscle is one of the four main muscles of mastication (chewing). It is attached to the ramus of the mandible and helps in jaw movements such as closing and chewing. When the masseter muscle contracts bilaterally, it elevates the mandible and closes the jaw (Fig. 1). In addition to this function, the masseter also causes protrusion, retraction, and side-to-side movement of the jaw, which helps in biting and chewing [1]. During biting and chewing, the masticatory muscles undergo the highest functional load, while exerting large forces. It has been reported that the resultant forces produced by the masticatory muscles on dental arches during maximum clenching range from 246.9 N to 2091.9 N in healthy humans [2]. Furthermore, the measurement of the mechanical characterization of the masseter has many applications in the medical field, including prediction of temporomandibular disorders [3] and bruxism [4], prediction of the aesthetic and functional consequences of orthognathic surgery [5] and predictive simulation and planning of cranio-maxillofacial surgery [6]. In recent years, robot assisted systems have been developed for mandible reconstruction and orthognathic surgery. Oral-rehabilitation robot was developed to provide robot-based massage therapy for temporomandibular joint disorders [7]. For instance, Ariji et al. [3] studied the changes in the mechanical behaviour of the masseter muscle in patients with myofascial pain after the robotic massage therapy. Furthermore, computational models were developed for masticatory system modelling and robotics, with applications in dental training, simulation of jaw movements, and speech therapy [8, 9, 10]. The development of these computational models offers significant advantages, including cost-effective and patient-specific treatment, and also reduces the need of specimens for testing while enhancing the better understanding of complex biomechanical processes. However, without the sufficient mechanical characterization of related tissues, masticatory modelling and computational models cannot be developed. Therefore, it is essential to study the mechanical characteristics of the masseter muscle, which is primarily involved in mastication. The studies on the mechanical behaviour of human facial soft tissues in regions including the oral, cheek, periorbital, forehead, jaw, nasal, temporal and auricular areas indicate that these tissues generally exhibit nonlinear, viscoelastic [11, 12, 13, 14, 15, 16, 17, 18] and anisotropic [19, 20, 21, 22, 23] characteristics. The existing in vivo studies on masseter muscle include, the functional and mechanical behaviour of masseter using imaging techniques such as sonography and ultrasound [24, 3, 25]; measurement of the mean stiffness of masseter muscle using shear wave elastography, with noticeable increase in stiffness of muscle indicating pathological tissues [26, 27]; study on the sliding behaviour and mechanical interaction between superficial musculoaponeurotic system (SMAS), subcutaneous and masseter muscle [28] using ultrasonography. The ex vivo studies involving the masseter muscle are limited. Zhuang et al. [29] studied the insertion and cutting forces of soft tissues of cadaver including masseter with the aim of improving accuracy in haptic feedback for virtual surgery system. The study characterized tissue response during insertion and cutting processes rather than intrinsic mechanical properties under tensile loading. Joy et al. [30] measured the stiffness of the masseter muscle and other soft tissues in Thiel soft-embalmed cadavers using shear wave elastography. Their study was conducted to demonstrate the effects of different embalming techniques for surgical training and anesthetists and to measure the tissue stiffness. Picard et al. conducted ex vivo tensile tests on human facial tissues including skin, hypodermis, zygomaticus muscle and masseter muscle, and analyzed their stress-strain responses, with the objective of developing biomechanical models of human facial soft tissue for applications such as surgical assistance and speech production. To characterize the history-dependent, viscoelastic and nonlinear mechanical behaviour of soft tissues, cyclic tensile testing method is commonly employed in which the tissues are subjected to several cycles of repeated loading and unloading at the same maximum stress and stress-strain characteristics are recorded. The soft tissues are subjected to repeated loading-unloading cycles until a stabilized stress-strain response is achieved. This process, referred to as preconditioning, is necessary to obtain a consistent and repeatable mechanical response [32]. The repeated loading-unloading during preconditioning leads to a history-dependent loss of stiffness, referred to as stress-softening in soft tissues and is termed as the Mullins effect in soft polymeric materials [32, 33].

**Fig. 1:**
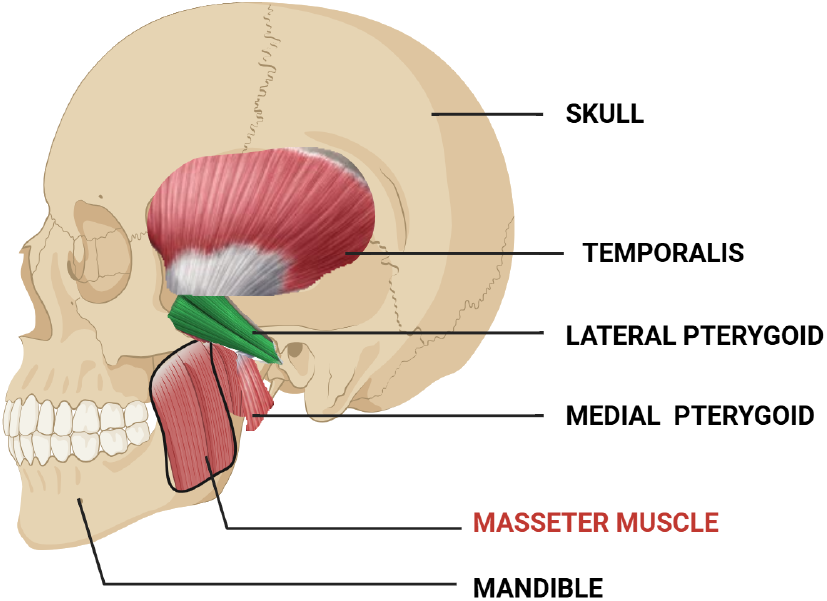
Schematic lateral view of the skull showing the anatomical location of the muscles of mastication, including the masseter, temporalis, medial pterygoid, and lateral pterygoid muscles, with emphasis on the masseter muscle.

To date, to be the best of the authors’ knowledge, there are no ex vivo studies that have employed cyclic tensile testing on the human masseter muscle. The existing tensile tests on the masseter muscle were conducted under monotonic loading conditions rather than cyclic loading [31]. Also, these studies did not capture the intrinsic mechanical properties of the masseter muscle required for a better understanding of the mastication biomechanics system.

The present work addresses this gap by performing controlled ex vivo cyclic tensile testing of the human masseter muscle under cyclic loading at increasing stretch levels, and the resulting stress-stretch response of the samples were investigated. The experimental data obtained from this study can be used for biomechanical modelling of the masseter and for the development and validation of constitutive models of the human masseter. Human cadavers are commonly used as biomechanical models and are commonly employed in ex vivo mechanical characterization. Specimens from fresh cadavers are suitable for conducting ex vivo tests, as they closely reproduce the natural mechanical properties. However, the availability of fresh cadavers is limited and therefore both preserved and fresh cadavers are used in ex vivo studies, even though the mechanical properties of fresh and preserved cadavers may differ [34]. Therefore, in this study, the effect of cadaver preservation on the mechanical behaviour of the masseter muscle is also investigated. The manuscript is organized as follows: The sample extraction, preparation and experimental methods are described in Section 2. In Section 3, the results of the experiments conducted are presented. The analysis on the time-dependent and viscoelastic behaviour is discussed in Section 4. Finally, in Section 5, the findings of the study are summarized along with suggestions for future works.

## 2. Materials and Methods

The anatomy of the human masseter muscle is described first, followed by the procedures for sample extraction and preparation, and the experimental setup.

### 2.1. Human masseter muscle

#### 2.1.1. Anatomy of the human masseter muscle

Masseter is one of the muscles of mastication and is quadrangular in shape. It has two parts: superficial and deep. The superficial part of the muscle originates from the temporal process of the zygomatic bone and the anterior part of the inferior border of the zygomatic arch, while the deep part originates from the zygomatic arch. The superficial masseter fibers run inferoposteriorly over the deep part and insert at the mandibular angle and the lateral surface of the mandibular ramus. The deep masseter fibers run inferiorly and insert along the mandibular ramus, superior to the region of insertion of the superficial portion. Fig. 1 shows a schematic lateral view of the human skull illustrating the anatomical location of the muscles of mastication, including the masseter, temporalis, medial pterygoid, and lateral pterygoid muscles. In this study, the masseter muscle is treated as a bulk tissue. To maintain consistency with the experimental approach, the masseter muscle is shown as a single anatomical entity.

The four main muscles of mastication include the masseter, temporalis, medial pterygoid and lateral pterygoid, all of which act on the mandible to produce jaw movements during opening and closing of the mouth. Among these, the masseter is the most powerful muscle in causing elevation and protrusion of the mandible by muscle contraction. Similar to the masseter muscle, contraction of the temporalis and medial pterygoid muscles also cause mandibular elevation, with temporalis additionally causing retraction and the medial pterygoid assisting in protrusion, while contraction of the lateral pterygoid facilitates protraction and depression of the mandible [1].

#### 2.1.2. Specimen extraction

Human masseter muscle tissue was obtained from nine cadaveric donors (six males and three females), aged between 78 and 98 years, for this study. The specimens comprised five fresh (unembalmed) cadavers and four formalin-embalmed. For embalming, a formalin solution was injected into the carotid artery and subsequently drained through the jugular vein. All donors were provided by the Laboratoire d’Anatomie des Alpes Françaises at Grenoble Alpes University, France. Dissections were performed in accordance with French regulations on postmortem testing, and were approved the University’s Ethics, Scientific, and Educational Committee. The fresh cadavers were stored under refrigerated conditions at 4 °C for up to 32 h postmortem. Dissections were performed within twenty-four hours of death to minimize tissue degradation and preserve the mechanical properties comparable to the in vivo state. Prior to incision, a hydro-dissection technique was employed to facilitate separation of facial tissue layers. Distilled water was injected into the subcutaneous tissue to preserve the hydration in the tissue (Fig. 2 (a)), to create a separation between the subcutaneous fat and the underlying muscle. This allowed removal of the fat without damaging the underlying muscle fibers. Following removal of the fat, any residual fluid was carefully eliminated to ensure that the specimen dimensions were not altered. A preauricular and submandibular incision was made. The incision extended from the posterior border of the mandible towards the mandibular angle. It was then continued anteriorly parallel to the inferior margin of the zygomatic arch. The skin and superficial fascia were carefully lifted and retracted, the subcutaneous fat layer was removed, and the masseter muscle was subsequently exposed, and isolated by dissection (Fig. 2(b).

**Fig. 2:**
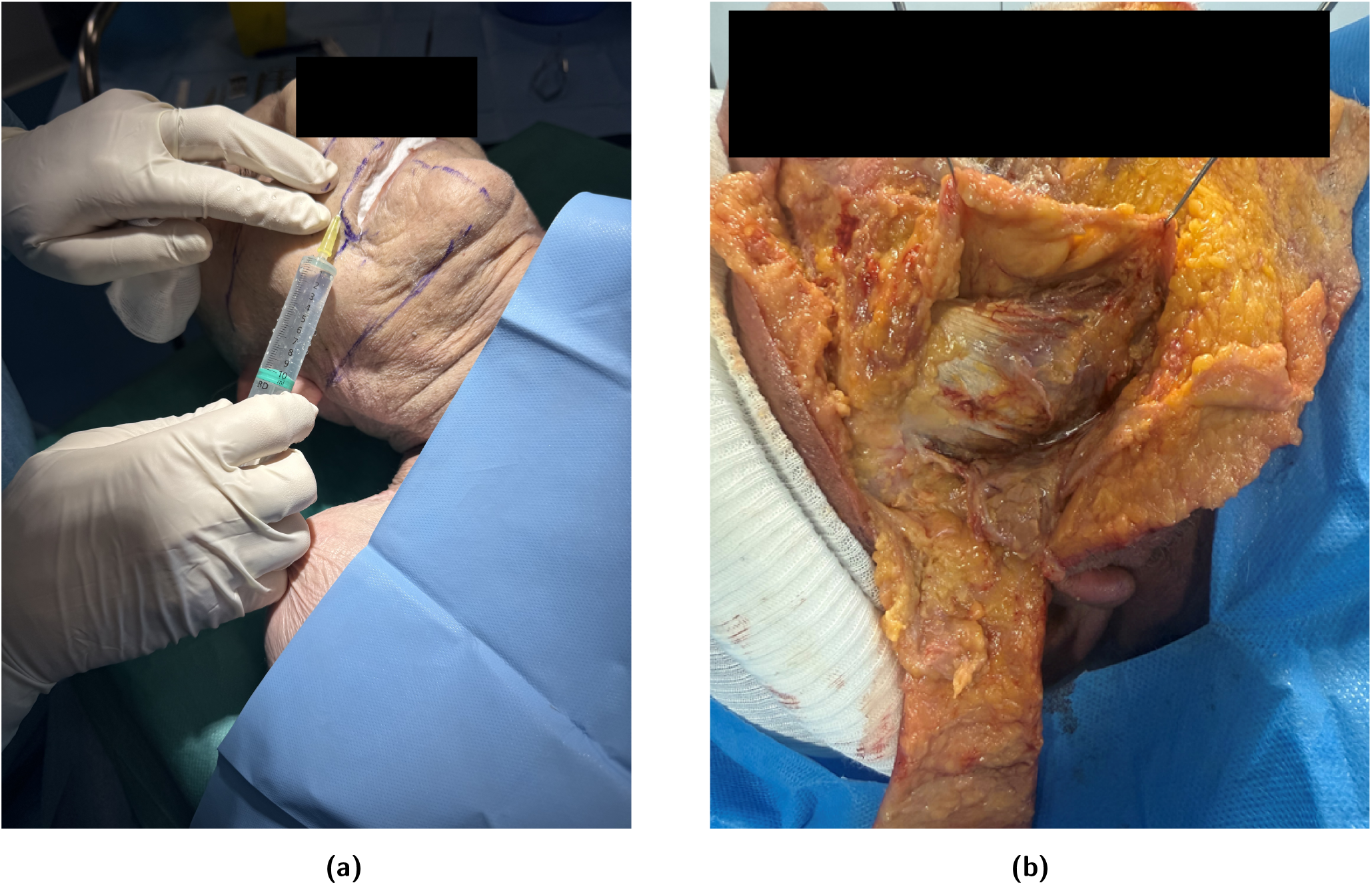
(a) Hydro-dissection of the facial subcutaneous tissue prior to incision (b) cadaveric dissection showing exposure of the masseter muscle

#### 2.1.3. Sample preparation

The specimens were cleaned with a scalpel and residual connective tissue was removed. An example of the masseter muscle specimen after isolation from a cadaver is shown in Fig. 3. The cleaned bulk tissue obtained from each cadaver was divided longitudinally into multiple specimens of approximately equal length. The number of specimens obtained from each cadaver varied depending on the size of the isolated bulk tissue.

**Fig. 3:**
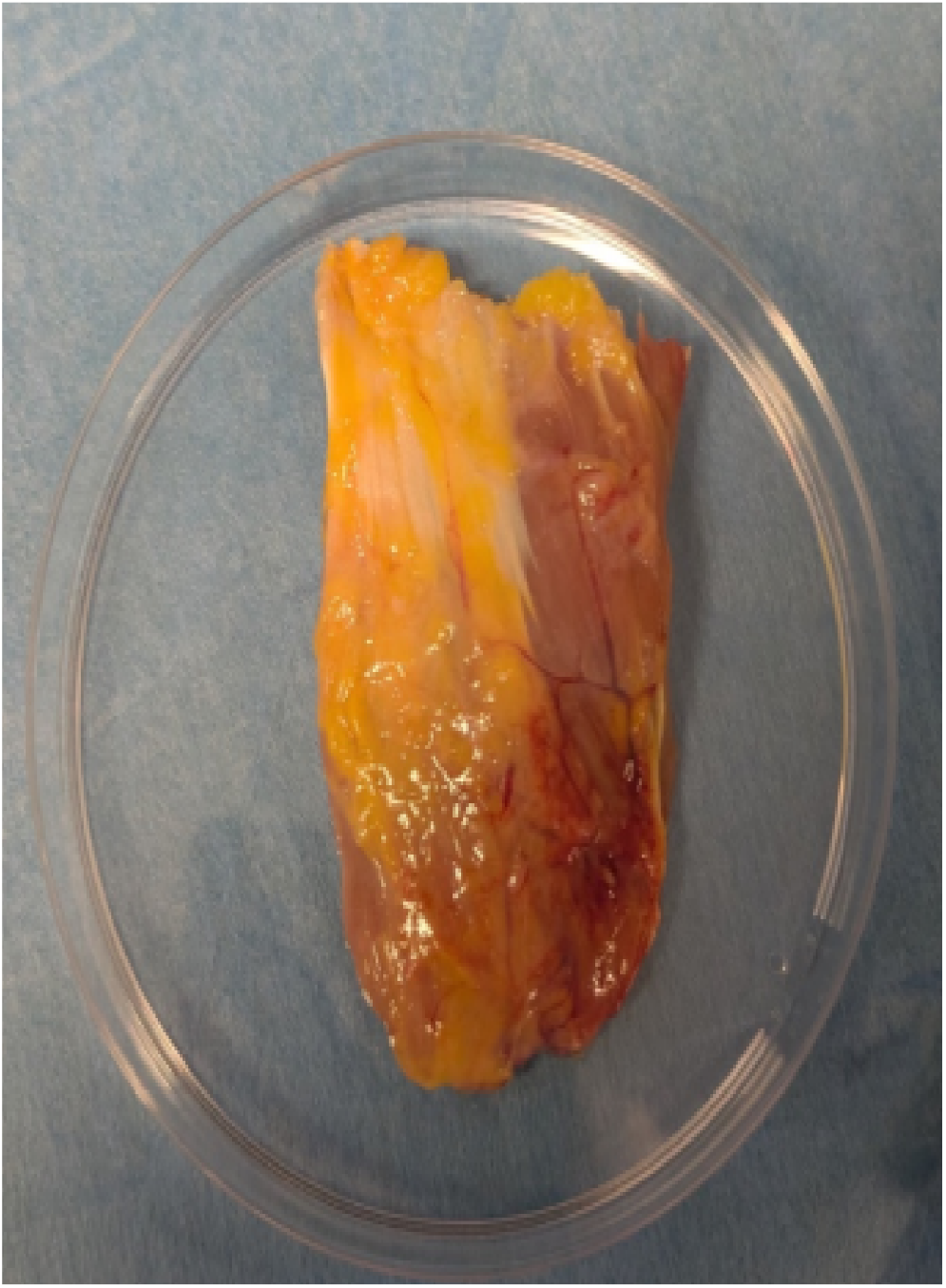
Masseter muscle bulk tissue isolated from a cadaver

All mechanical tests were completed within 48 hours of dissection, during which the specimens were stored in physiological saline solution (0.9% NaCl) at 4 °C in a refrigerator.

### 2.2. Experimental setup

#### 2.2.1. Uniaxial tensile testing

Before loading the masseter sample within the tensile testing machine, it was secured within the grips using a special device, as shown in Fig. 4(a). To mount the tissue sample within the grips, the sample was first placed on the base grips and the back support of the device. Next, the sample was flattened and aligned centrally. The upper grips were placed over the base grips, and the screws were tightened using a torque limiter set to 0.5 Nm to prevent sample slipping during testing and ensure consistency between tests. The longer screws on either side of the grips were then tightened to complete the assembly. The assembled unit was mounted within the mechanical testing system (MTS Criterion C41, MTS Systems, Minnesota), by fixing its lower part to the machine base and its upper part to a highly sensitive 25 N load cell, as shown in Fig. 4(b). Once the sample was secured, the side screws were loosened to release the assembly and the back support was removed. The final experimental configuration before the commencement of the tests is shown in Fig. 4(c). At this stage, the width and thickness of the samples were measured at three points along the length using calipers, and an average value was used for subsequent analysis. Uniaxial tensile testing was performed by applying deformation through upward traction of the upper grip attached to the crosshead and load cell, while the lower grip remained fixed. Sample deformation was obtained using an extensometer installed in the testing system, which recorded the crosshead displacement. Strain was then calculated from the grip-to-grip length of the sample. The sample geometry was maintained with a length-to-width ratio of approximately 4:1, consistent with ASTM standards for uniaxial tensile testing (ASTM, 2013). The testing system was operated using MTS TestSuite software, through which all test parameters were defined.

**Fig. 4:**
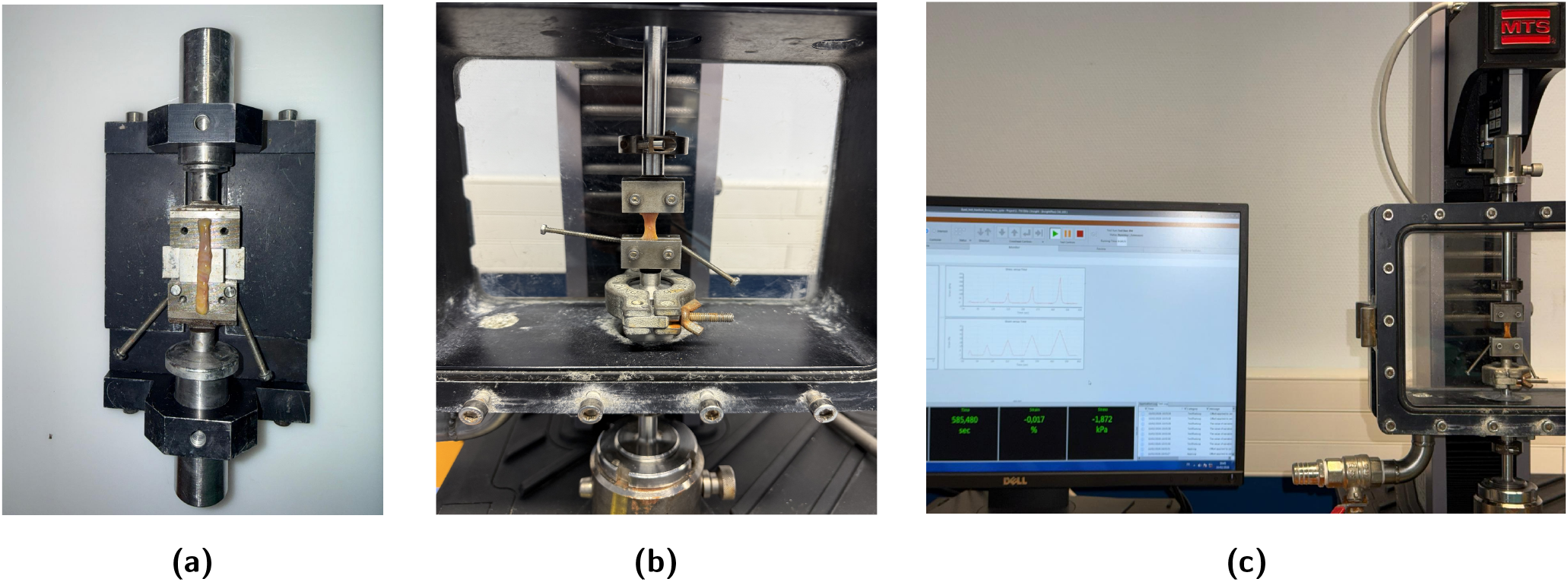
Experimental mounting and tensile testing configuration of the masseter specimen: (a) Sample positioned within the gripper; (b) Sample loaded in the machine; (c) Final experimental configuration prior to stretch-controlled loading.

#### 2.2.2. Cyclic tests

In this study, cyclic tests were conducted to investigate the viscoelastic behaviour of the masseter muscle. The cyclic tests comprised two protocols, namely the continuous stretch level cyclic loading (continuous stretch protocol) and the constant stretch level cyclic loading (constant stretch protocol). Under the continuous stretch protocol, the cyclic tests comprised a series of loading and unloading phases with increasing stretch levels of 1.05, 1.10, 1.15, 1.20, and 1.25, and the stress response of the tissue was continuously recorded. Each cycle consisted of loading to the prescribed stretch level, followed by unloading and a 60-second hold period. The stretch-time curve of this testing protocol can be seen in Fig. 5(a). The maximum stretch level of 1.25 and a stretch increment of 5% were selected to avoid premature tissue rupture while capturing the nonlinear viscoelastic behaviour of the tissue. If the tissue did not rupture at the maximum stretch level of 1.25 under the continuous stretch protocol, the sample was subsequently stretched beyond 1.25 until failure, and the corresponding rupture stress and stretch were recorded. The strain rate-dependent behaviour of the tissue was investigated by conducting cyclic tests at two different strain rates, 0.01 % s^−1^ and 0.1 % s^−1^. Only one cycle was applied at each stretch level in this testing protocol. Considering the availability of a larger number of specimens (128 tests), independent tests were prioritised to enable improved inter-group comparisons (fresh vs formalin; 0.01 % s^−1^ vs 0.1 % s^−1^) and robust statistical analysis. In addition to characterising the viscoelastic behaviour, stress-softening, residual strain, hysteresis, and rupture behaviour were examined. Under the constant stretch protocol, a cycle consisting of loading–unloading followed by a 60 s hold was repeated five times at a maximum stretch level of 1.2, and the responses were recorded. The stretch-time curve of this testing protocol is shown in Fig. 5(b). All experiments were conducted on both fresh and formalin-embalmed tissue samples to assess the effect of tissue preservation. Similar to the continuous stretch protocol, the effect of conditioning (stress-softening), as well as residual strain and hysteresis was also examined. Each sample was tested only once, and no specimen was subjected to more than one complete test. All experiments were conducted at ambient temperature.

**Fig. 5:**
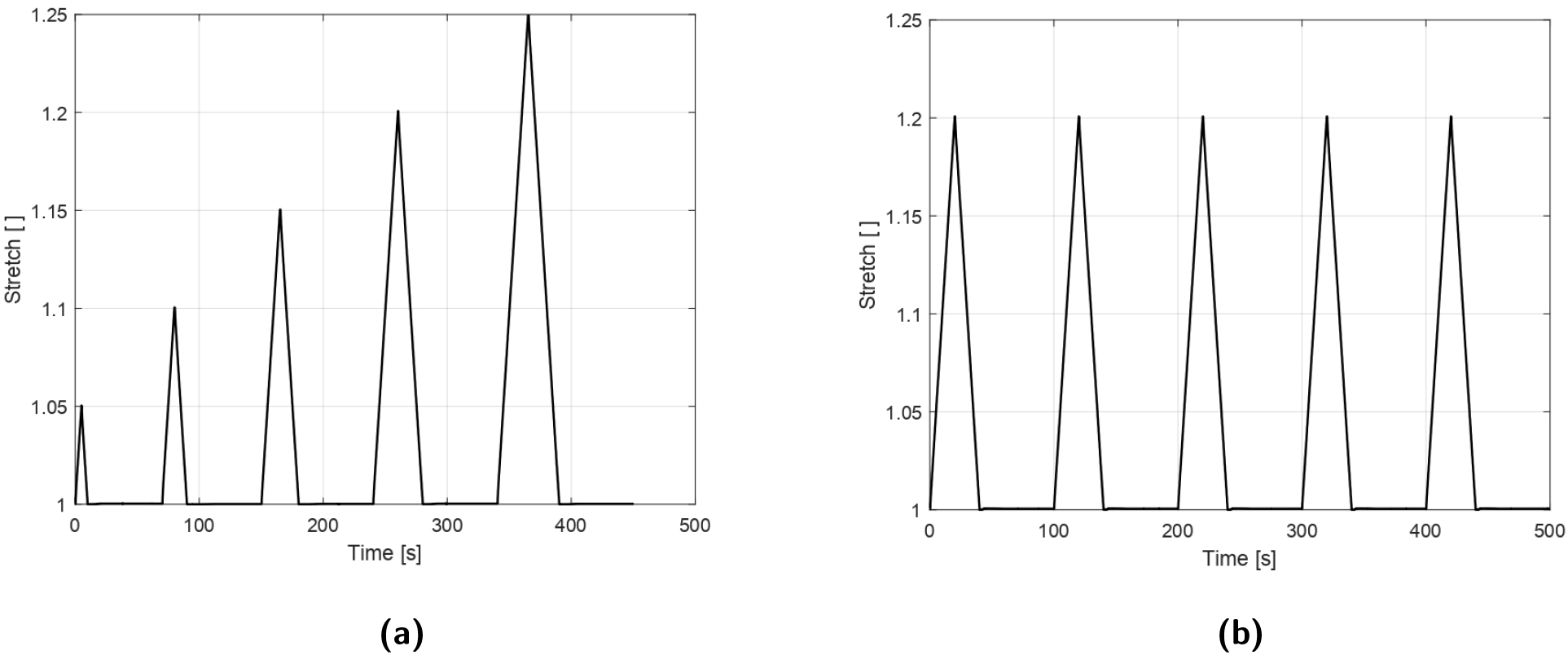
Stretch–time curves of the mechanical test protocols: (a) cyclic loading protocol with increasing stretch levels (1.00–1.25) and 60 s hold period at each level; (b) Repeated cyclic loading protocol at constant maximum stretch level (*λ* = 1.2) for five cycles

#### 2.2.3. Definition of mechanical quantities

In this study, uniaxial tensile testing was conducted under stretch controlled conditions. The measurements of force and crosshead displacement were continuously recorded, and the corresponding stress-stretch values were calculated using the following definitions: The stretch *λ*, is defined as:

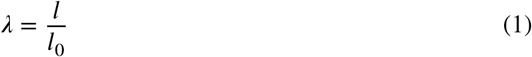

where *l* and *l*_0_ are the instantaneous and initial lengths of the sample, respectively. The nominal strain is calculated as:

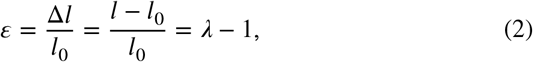

where Δ*l* is the crosshead displacement used to estimate sample deformation.

The stress is expressed as nominal stress (first Piola– Kirchhoff stress) in N/mm^2^ (MPa), defined as:

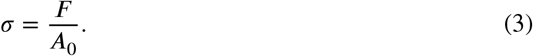

where *F* is the applied tensile force recorded by the load cell and *A*_0_ is the original undeformed cross-sectional area.

### 2.3. Donor demographics and specimen distribution

The demographics of the nine cadavers used for the continuous stretch protocol study and their preservation condition are presented in Table 1 and the demographics of the three cadavers used for the constant stretch protocol are presented in Table 3. The total number of specimens tested per test condition (i.e. preservation condition and strain rate) for each cadaver tested under continuous stretch protocol, along with their geometric dimensions (mean± Standard Deviation (SD)), is provided in Table 2 and the total number of speciments tested under constant stretch protocol is given in Table 4. Due to the variability of biological tissue, specimen dimensions such as width and thickness varied both between and within cadavers, as shown in Table 2.

**Table 1.** Demographic characteristics and preservation status of the cadaveric donors (n = 9).

| Cadaver | Preservation | Sex | Age (years) | Weight (kg) | Height (cm) |
| --- | --- | --- | --- | --- | --- |
| 1 | Formalin | Male | 78 | 70 | 175 |
| 2 | Formalin | Female | 98 | 60 | 150 |
| 3 | Formalin | Male | 79 | 85 | 175 |
| 4 | Formalin | Male | 86 | 60 | 160 |
| 5 | Fresh | Male | 85 | 70 | 170 |
| 6 | Fresh | Female | 95 | 55 | 150 |
| 7 | Fresh | Female | 89 | 60 | 160 |
| 8 | Fresh | Male | 81 | 70 | 160 |
| 9 | Fresh | Male | 84 | 75 | 175 |

**Table 2.** Cadaver-wise specimen counts tested under different preservation conditions and strain rates, with geometric dimensions (mean ± SD)

| Cadaver preservation | Strain rate | Cadaver | No. of specimens | Total | Length (mm) | Width (mm) | Thickness (mm) |
| --- | --- | --- | --- | --- | --- | --- | --- |
| Fresh | 0.01 s<sup>-1</sup> | 1 | 6 | 23 | 34.0 ± 1.2 | 4.0 ± 0.5 | 1.4 ± 0.3 |
|  |  | 2 | 6 |  |  |  |  |
|  |  | 3 | 5 |  |  |  |  |
|  |  | 4 | 3 |  |  |  |  |
|  |  | 5 | 3 |  |  |  |  |
| | 0.1 s<sup>-1</sup> | 1 | 6 | 27 | 35.2 ± 1.0 | 4.1 ± 0.4 | 1.5 ± 0.4 |
|  |  | 2 | 6 |  |  |  |  |
|  |  | 3 | 6 |  |  |  |  |
|  |  | 4 | 6 |  |  |  |  |
|  |  | 5 | 3 |  |  |  |  |
| Formalin | 0.01 s<sup>-1</sup> | 1 | 15 | 44 | 31.6 ± 1.5 | 3.8 ± 0.6 | 1.7 ± 0.5 |
|  |  | 2 | 7 |  |  |  |  |
|  |  | 3 | 11 |  |  |  |  |
|  |  | 4 | 11 |  |  |  |  |
| | 0.1 s<sup>-1</sup> | 1 | 10 | 34 | 33.8 ± 1.3 | 3.9 ± 0.5 | 1.3 ± 0.4 |
|  |  | 2 | 14 |  |  |  |  |
|  |  | 3 | 10 |  |  |  |  |

**Table 3.**
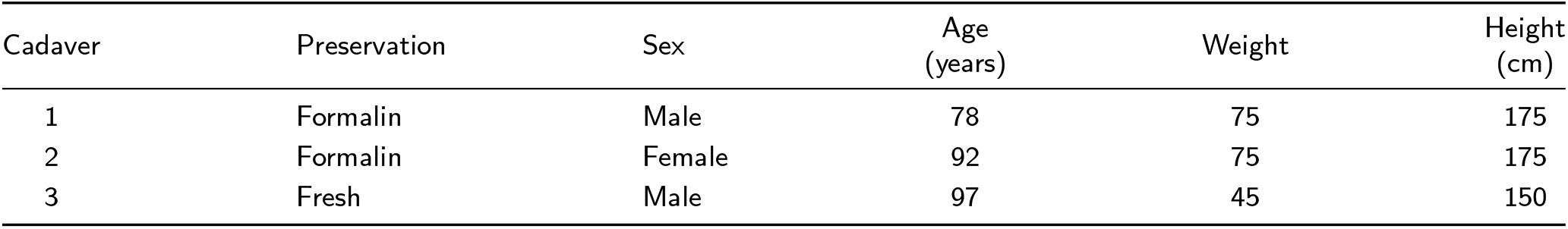
Demographic characteristics and preservation status of the cadaveric donors used for the constant stretch protocol (n = 3).

**Table 4.** Cadaver-wise specimen counts tested under different preservation conditions and strain rates for the constant stretch protocol, with geometric dimensions (mean ± SD).

| Cadaver preservation | Strain rate | Cadaver | No. of specimens | Total | Length (mm) | Width (mm) | Thickness (mm) |
| --- | --- | --- | --- | --- | --- | --- | --- |
| Fresh | 0.01 s<sup>-1</sup> | 1 | 2 | 4 | 31.1 ± 1.2 | 4.3 ± 0.5 | 1.2 ± 0.3 |
|  |  | 2 | 1 |  |  |  |  |
|  |  | 3 | 1 |  |  |  |  |
| | 0.1 s<sup>-1</sup> | 1 | 1 | 3 | 33.4 ± 1.1 | 4.1 ± 0.4 | 1.4 ± 0.4 |
|  |  | 2 | 1 |  |  |  |  |
|  |  | 3 | 1 |  |  |  |  |
| Formalin | 0.01 s<sup>-1</sup> | 1 | 1 | 3 | 31.2 ± 1.3 | 4.6 ± 0.4 | 1.1 ± 0.2 |
|  |  | 2 | 1 |  |  |  |  |
|  |  | 3 | 1 |  |  |  |  |
| | 0.1 s<sup>-1</sup> | 1 | 1 | 4 | 33.2 ± 1.3 | 4.8 ± 0.3 | 1.6 ± 0.4 |
|  |  | 2 | 2 |  |  |  |  |
|  |  | 3 | 1 |  |  |  |  |

### 2.4. Statistical method

#### Continuous stretch protocol

The specimens were grouped according to preservation condition (fresh or formalin) and were tested either at 0.01 s^−1^ or 0.1 s^−1^ strain rate. Due to the variability observed in the response of the samples within the same test condition, a statistical approach was applied to identify the most representative stress–stretch response within each test condition for subsequent analysis. For each test condition, the tangent modulus, *E*_tan_, was calculated from the nominal stress–stretch curves of individual tensile tests. The tangent modulus of each tensile test was computed from the first loading curve of the stress–stretch response between the stretch levels of 1.00 and 1.05. The tangent modulus within this interval was obtained by averaging the pointwise tangent moduli calculated at discrete points within this range. The tangent modulus at a point on the loading curve is defined by:

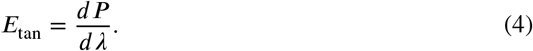

In this study, the tangent modulus at a point *i* was determined numerically using central difference approximation. After the determination of tangent modulus, a histogram with 20 bins was plotted for each test condition with tangent modulus on the x-axis and density on the y-axis. To assess normality, the following null hypothesis was formulated:

*“For each test condition, the tangent modulus follows a normal distribution.”* A Shapiro-Wilk test was conducted using Python, with a significance level of *α* = 0.05. The null hypothesis was rejected if *p* < 0.05. If the data were normally distributed, the mean value was used as the representative measure. If the data were not normally distributed, a goodness-of-fit analysis using Kolmogorov–Smirnov (KS) test was carried out to assess whether the data followed a Fréchet distribution [35]. For this test, the null hypothesis was: *“The tangent modulus follows a Fréchet distribution.”* The test was conducted at *α* = 0.05, and the null hypothesis was retained if *p* > 0.05. In this case, mode of the fitted Fréchet distribution was used as the representative value. The Fréchet distribution was used in this study, as the tangent modulus of all tensile tests exhibited a right skewed distribution and also and the Fréchet distribution is suitable for the characterization of soft biological tissue properties [36]. Additionally, due to the non-normality of the data, statistical comparisons between different test conditions were performed using the non-parametric Mann– Whitney U test with a significance level of *p* < 0.05. These comparisons were conducted on the quantitative parameters derived from the experimental data, including tangent modulus, energy dissipation (hysteresis), stress-softening index, residual strain, and rupture stress and stretch.

#### Constant stretch protocol

Similar to continuous stretch protocol, the specimens were grouped according to preservation condition (fresh or formalin) and were tested at strain rates of 0.01 s^−1^ or 0.1 s^−1^. However, the number of specimens tested under the constant stretch protocol was limited. Since normality tests are known to have low statistical power for small sample sizes and may fail to detect deviations from normality [37], the representative curve for each test condition was selected using the median as it is less sensitive to outliers and skewed data. Consistent with the continuous stretch protocol, non-parametric statistical comparisons were performed on the experimentally derived parameters using the Mann– Whitney U test.

### 2.5. Hysteresis area and residual strain

Hysteresis area (*A*_*HA*_) was calculated as the difference between the area under the loading curve (*A*_*L*_) and the area under the unloading curve (*A*_*UL*_) [38]:

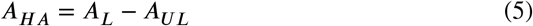

The area under the loading and unloading curves were determined as:

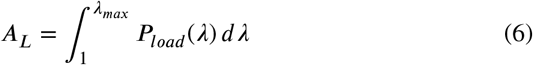

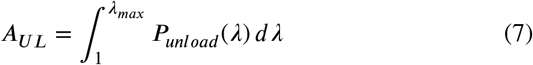

where *P*_*load*_ and *P*_*unload*_ are the nominal stresses of the loading and unloading curves, respectively.

The hysteresis was then normalized as:

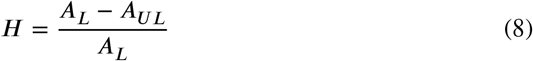

Residual strain, defined as *λ*_*r*_ − 1, is obtained from the unloading curve at the stretch where the stress becomes zero, and represents the permanent deformation of the tissue. [39].

In this study, hysteresis area and residual strain were calculated for each loading-unloading cycle at all five stretch levels. The determination of hysteresis area and the residual strain is illustrated in Fig. 6.

**Fig. 6:**
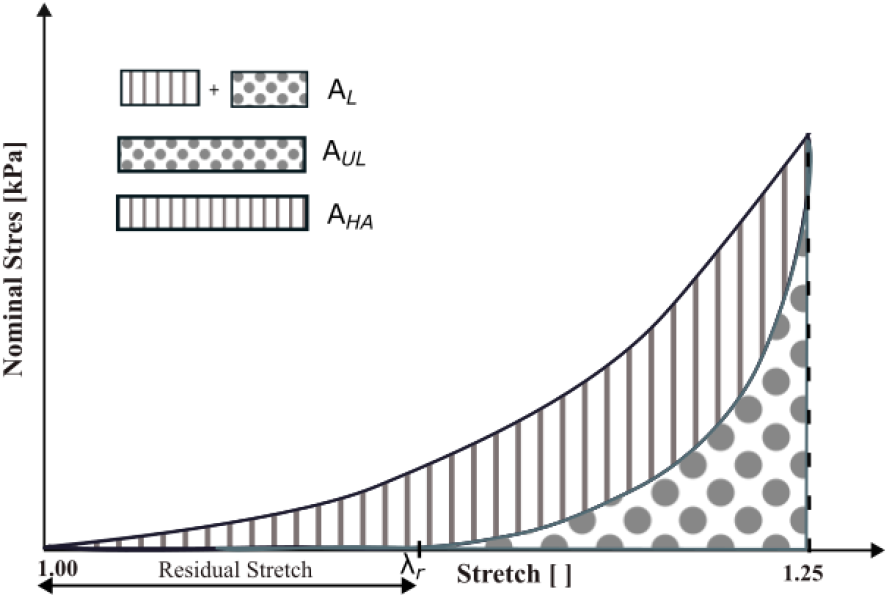
Schematic diagram illustrating hysteresis and residual stretch during cyclic loading-unloading to a maximum stretch level. *A*_*L*_ indicates the area under the loading curve, *A*_*UL*_ denotes the area under the unloading curve and *A*_*HA*_ represents the hysteresis area that corresponds to energy dissipated during the cycle. *λ*_*r*_ denotes the stretch at zero stress upon unloading

### 2.6. Stress-softening

When the tissue is subjected to cyclic loading, it exhibits reduced stiffness due to the mechanism of stress-softening. Stress-softening also known as Mullins effect, is characterized by a reduction in the stress in subsequent cycles when compared to initial loading cycle at the same stretch level [39, 40] and was determined by computing the normalized difference in strain energy density between the virgin envelope (*W*_*virgin*_) and the reloading curve (*W*_*reload*_) as follows:

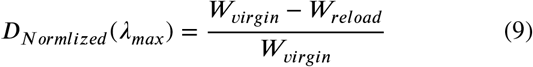

where *λ*_*max*_ is the maximum stretch level reached during the loading cycle.

The strain energy density of the virgin envelope and the reloading curve was calculated by computing the area under their corresponding stress-stretch curves as follows:

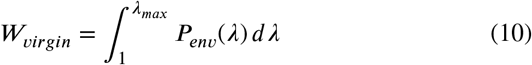

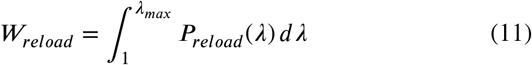

where *P*_*env*_ and *P*_*reload*_ denote the nominal stresses of the virgin envelope and the reloading curve.

In this study, stress-softening is investigated in two testing protocols: (i) continuous stretch level cyclic loading, which is the primary protocol of this study to investigate viscoelastic behaviour, and (ii) constant stretch level cyclic loading. Continuous stretch level cyclic loading refers to a gradual increase in stretch levels at each cycle, whereas constant stretch level cyclic loading refers to repeated cyclic loading at a fixed maximum stretch level. The schematics showing stress-softening are illustrated in Fig. 7 (a) and (b).

**Fig. 7:**
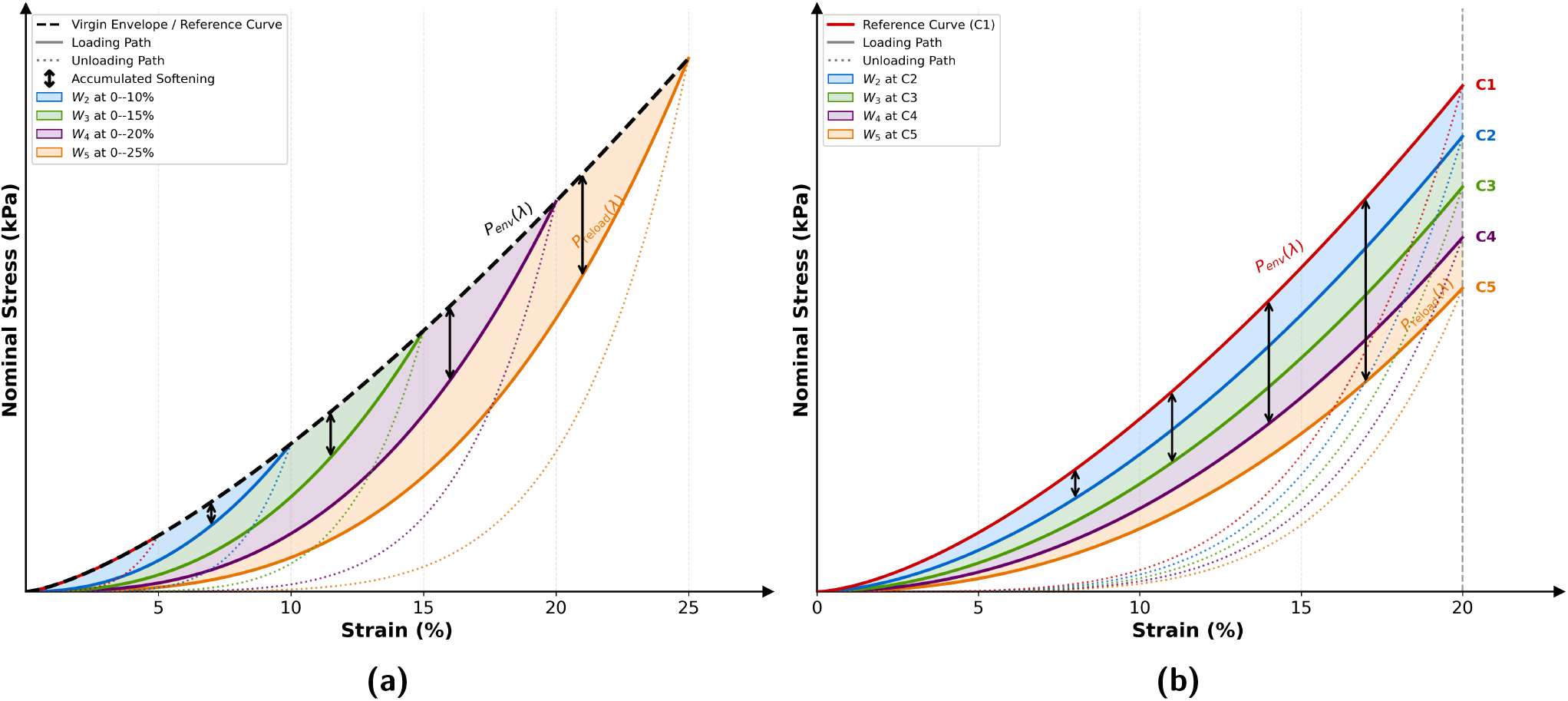
Schematics illustrating stress-softening: (a) Continuous stretch level cyclic loading, where the stretch levels are increased progressively (5–25%) in successive cycles, and the virgin envelope is constructed by connecting the peak stress values of the primary loading cycles. The shaded regions represent the accumulated stress-softening relative to the virgin envelope. (b) Constant stretch level cyclic loading, where repeated cyclic loading is applied at a fixed maximum stretch level. The incremental reduction in stress between successive cycles is shown from C_1_–C_5_. The shaded regions represent the accumulated softening with respect to the first loading cycle. The nominal stress corresponding to the virgin envelope is denoted by *P*_env_(*λ*), while the nominal stress associated with the reloading curves is denoted by *P*_reload_(*λ*). *P*_reload_(*λ*) is indicated on the final reloading curve, which is representative of the reloading response across cycles.

## 3. Experimental Results

### 3.1. Presentation of cyclic tests results and variability in tissue behaviour

For a specific test condition, a large variation in the response of the samples was observed. This can be seen in Fig. 8 (a), which depicts the rupture stress-stretch response of all fresh cadaver samples, while Fig. 8 (b) shows the corresponding response for formalin-preserved samples. Hence, to obtain the most representative stress-stretch data for each test condition, the statistical approach as described in Section 2.4. was employed. The results of this analysis are presented in the following section.

**Fig. 8:**
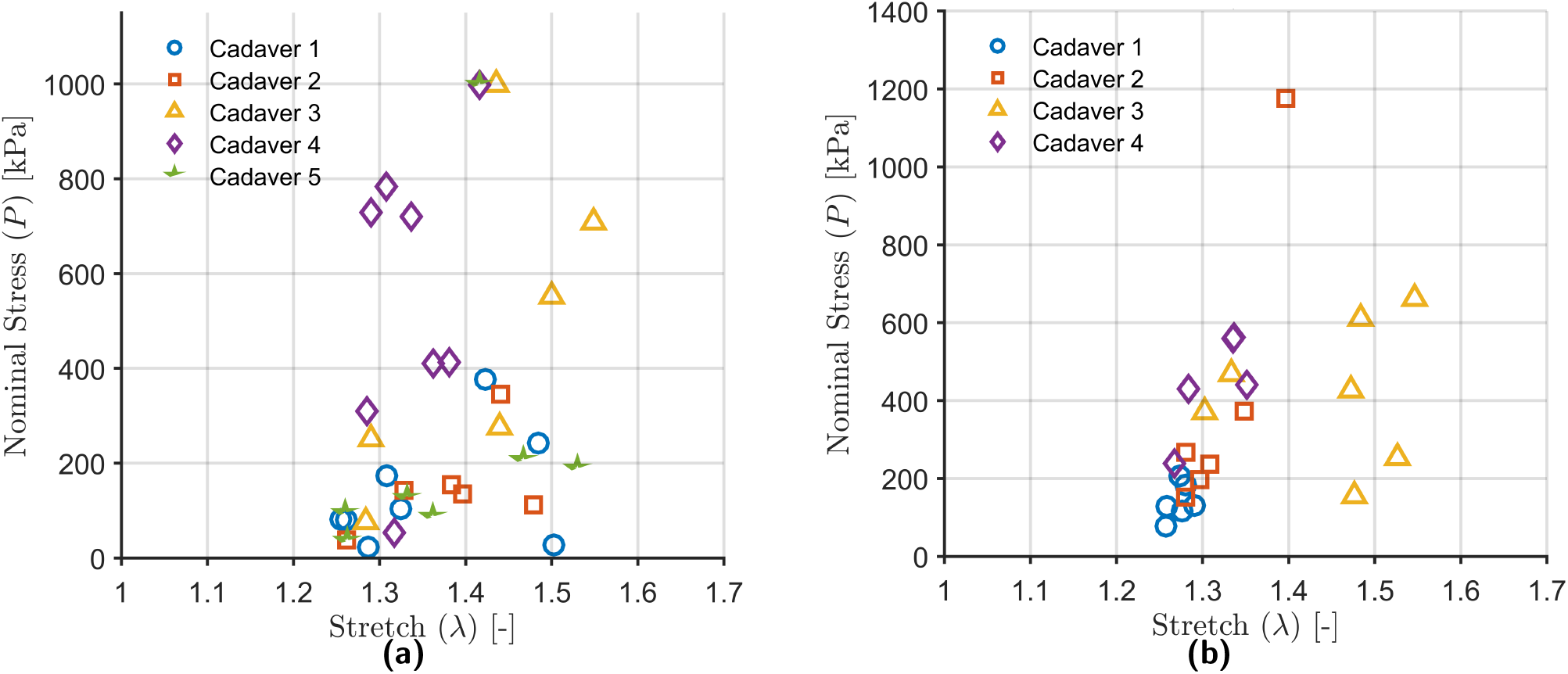
Rupture stress–stretch response of masseter muscle of fresh cadaver samples at 0.01 s^−1^ strain rate (a) and formalin-preserved cadaver samples 0.1 s^−1^ strain rate (b). Each marker represents an individual sample, highlighting the dispersion across samples and strain rates.

Although rupture behaviour was not evaluated under the constant stretch protocol, variability in the cyclic response of the specimens was observed across the test conditions in terms of parameters such as tangent modulus and energy dissipation. Due to the limited number of samples and the associated uncertainty in the underlying data distribution, the most representative response for each test condition was selected using the median of the tangent modulus, as described in Section 2.4.

### 3.2. Statistical analysis and representative curve selection

The histograms of the tangent modulus for each test condition are plotted as shown in Fig. 9 (a)–(d). All test conditions exhibited right-skewed histograms, indicating non-normal dispersion of the tangent moduli. This was confirmed by conducting the Shapiro–Wilk test, resulting in p values less than 0.05 for all test conditions. Thus, the null-hypothesis for normal distribution was rejected. For the non-normal distribution of data, a Fréchet distribution was fitted and evaluated using the KS test. The null hypothesis was not rejected as the p values were greater than 0.05. Hence, for each test condition, the stress-stretch curve whose tangent modulus is closest to the mode of the Fréchet distribution was chosen as the representative curve for further analysis. The statistical results for each test condition are summarized in Table 5. For the constant stretch protocol, the tangent modulus, *E*_tan_, was calculated in a manner similar to the continuous stretch protocol. However, the number of samples tested under each test condition was limited; therefore, unlike the continuous stretch protocol, the representative curve for each test condition was selected using the median of the tangent modulus, as described in Section 2.4

**Table 5.** Statistical analysis and representative curve selection for each test condition.

| Test condition | No. of tests, $n$ | Shapiro–Wilk p-value | Normality decision | KS p-value | Distribution selected | Fréchet mode (MPa) | Representative $E_{\text{tan}}$ (MPa) |
| --- | --- | --- | --- | --- | --- | --- | --- |
| $0.01 \text{ s}^{-1}$ fresh | 23 | 0.017 | Non-normal | 0.819 | Fréchet | 0.353 | 0.360 |
| $0.1 \text{ s}^{-1}$ fresh | 27 | 0.032 | Non-normal | 0.997 | Fréchet | 1.034 | 1.040 |
| $0.01 \text{ s}^{-1}$ formalin | 44 | 0.001 | Non-normal | 0.998 | Fréchet | 0.916 | 0.936 |
| $0.1 \text{ s}^{-1}$ formalin | 34 | $1.46 \times 10^{-7}$ | Non-normal | 0.549 | Fréchet | 2.030 | 2.101 |

**Fig. 9:**
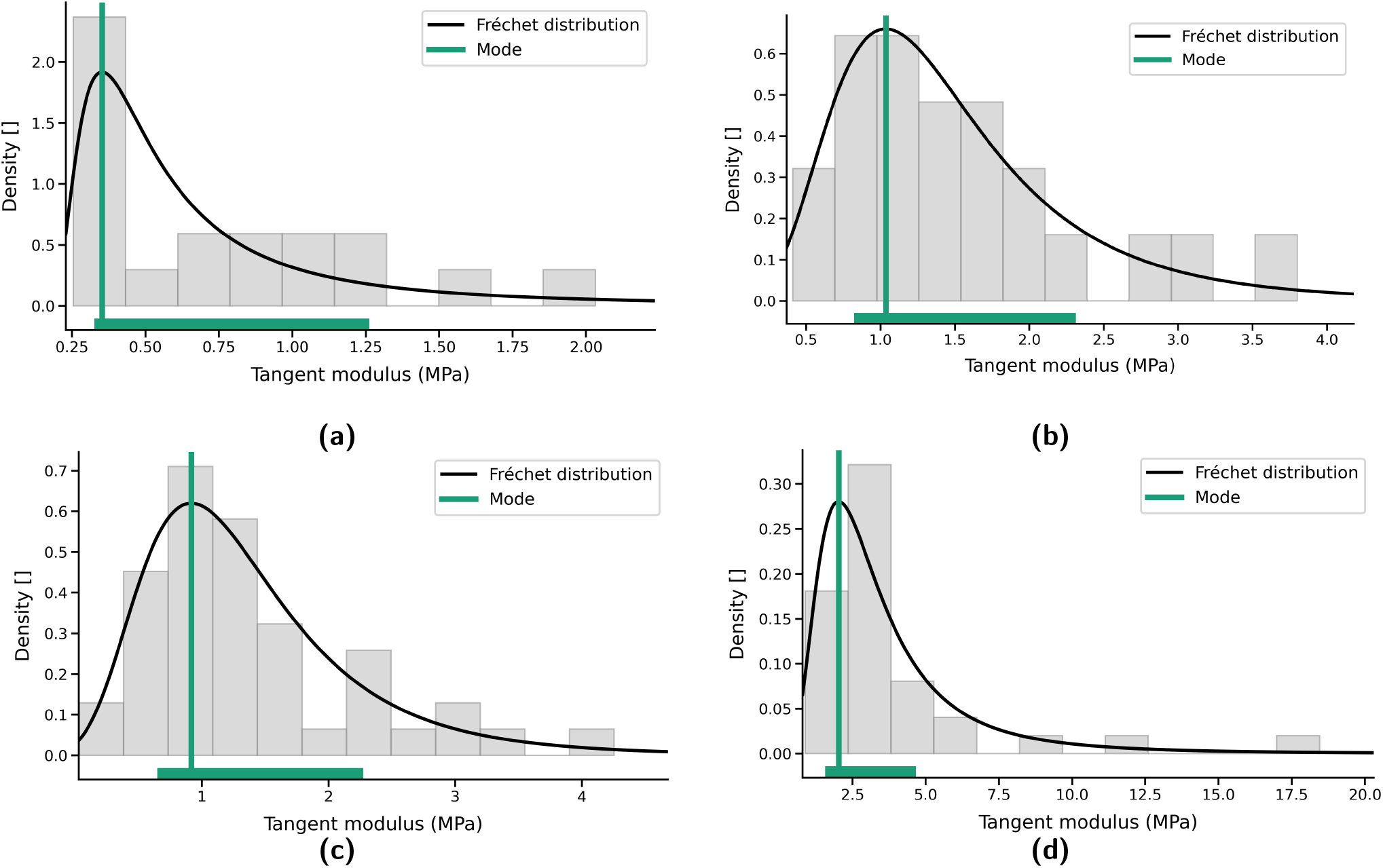
Histogram distributions of tangent modulus obtained from the stress–stretch response, together with the probability distribution graph for each test condition: (a) 0.01 s^−1^ fresh; (b) 0.1 s^−1^ fresh; (c) 0.01 s^−1^ formalin; and (d) 0.1 s^−1^ formalin. The vertical green line denotes the mode of the fitted distribution, while the horizontal green bar indicates the range containing 70% of the tangent moduli

### 3.3. Cyclic stress–stretch behaviour

The representative stress–stretch responses of the masseter muscle under both continuous stretch and constant stretch protocols are shown in Fig. 10–Fig. 11 and Fig. 12– Fig. 13, respectively. These figures illustrate the effect of strain rate and cadaver preservation on the mechanical response of the tissue. Under the continuous stretch protocol (Fig. 10 and Fig. 11), it can be observed that the stress– stretch curves exhibit a J-shaped response consisting of three regions [35]. The initial toe region and the final region show nonlinear behaviour, with a lower increase in stress at lower stretch levels in the toe region and a steeper increase in stress at higher stretch levels in the final region, while the intermediate region exhibits an approximately linear response. A noticeable increase in the slope of the stress– stretch curve is observed beyond the intermediate stretch levels. In contrast, under the constant stretch protocol (Fig. 12 and Fig. 13), the stress–stretch response is characterized by repeated loading–unloading cycles at a fixed maximum stretch. A progressive reduction in stress can be observed across the cycles.

**Fig. 10:**
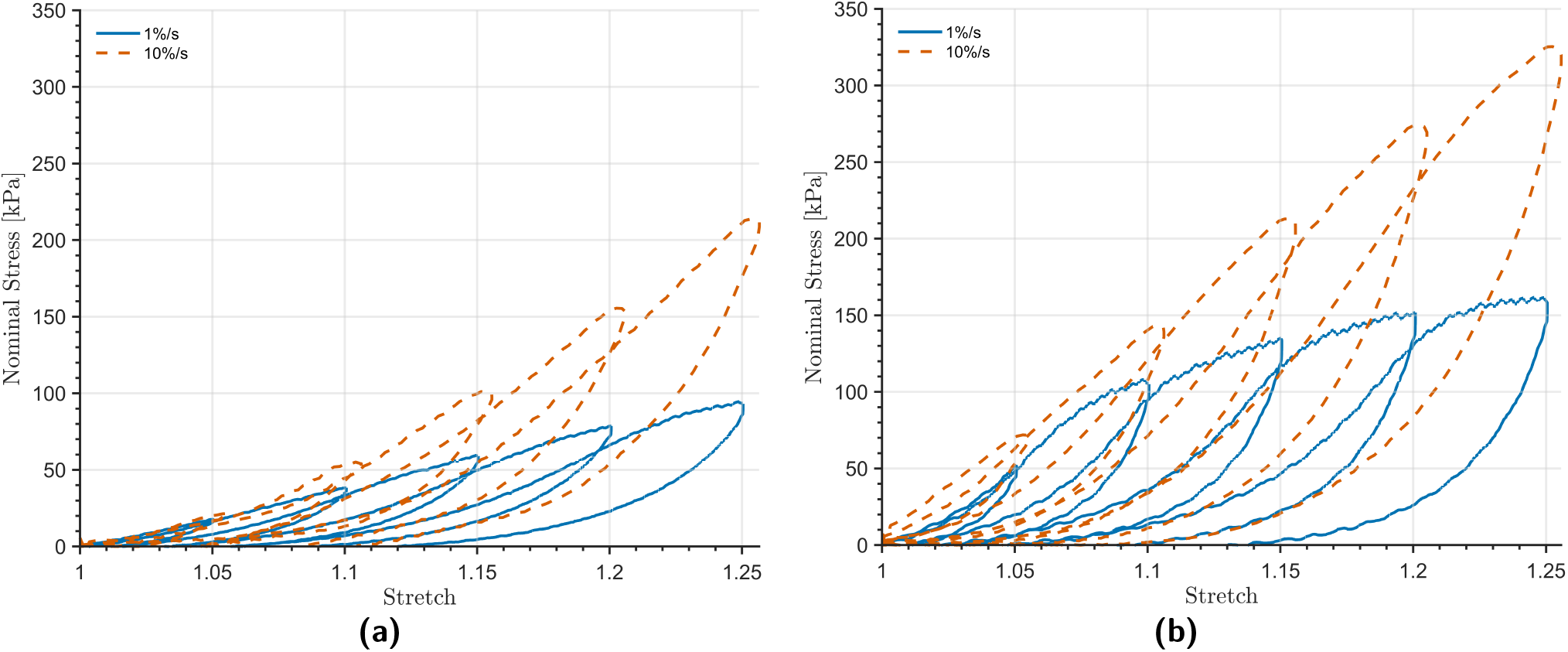
Representative stress–stretch responses of the masseter muscle samples under continuous stretch-level cyclic loading at two strain rates (0.01 s^−1^ and 0.1 s^−1^). (a) Fresh samples and (b) formalin-preserved samples.

**Fig. 11:**
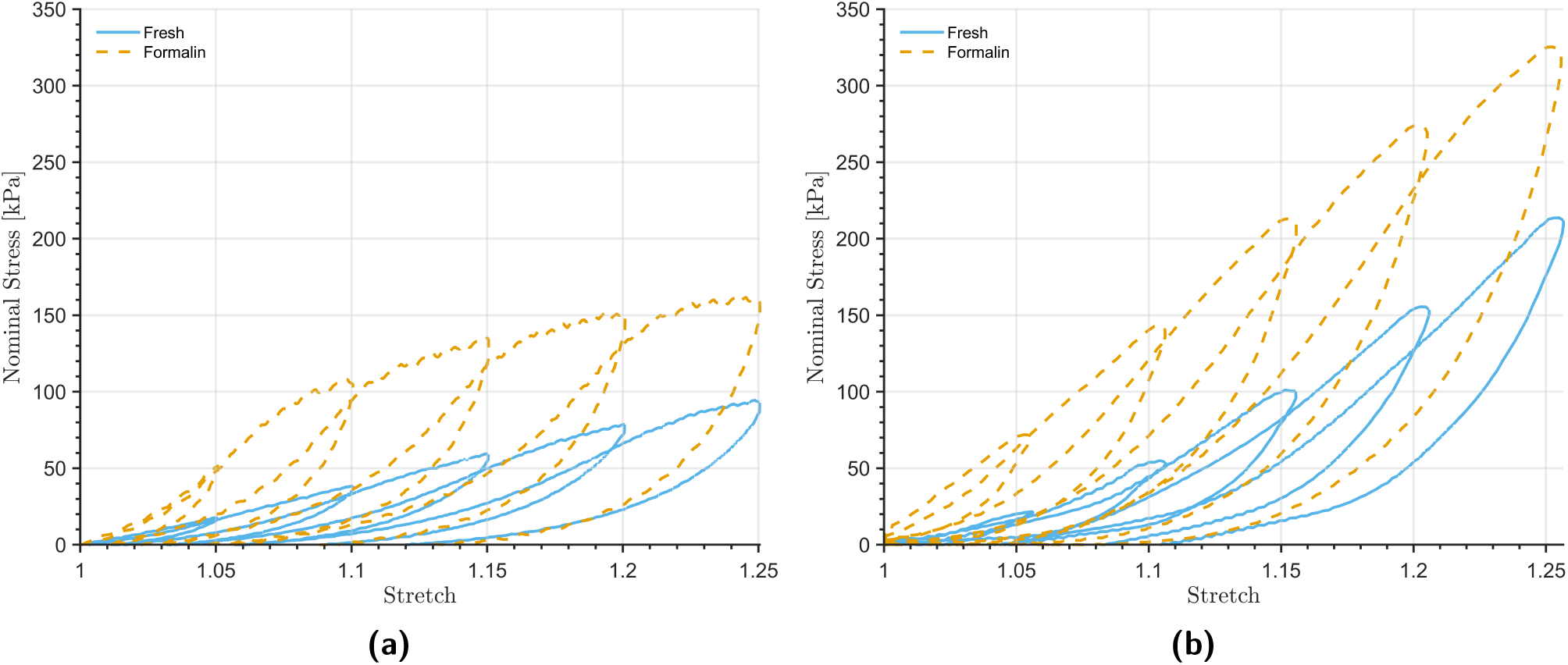
Representative stress–stretch responses of fresh and formalin-preserved masseter muscle samples under continuous stretch-level cyclic loading at (a) 0.01 s^−1^ and (b) 0.1 s^−1^

**Fig. 12:**
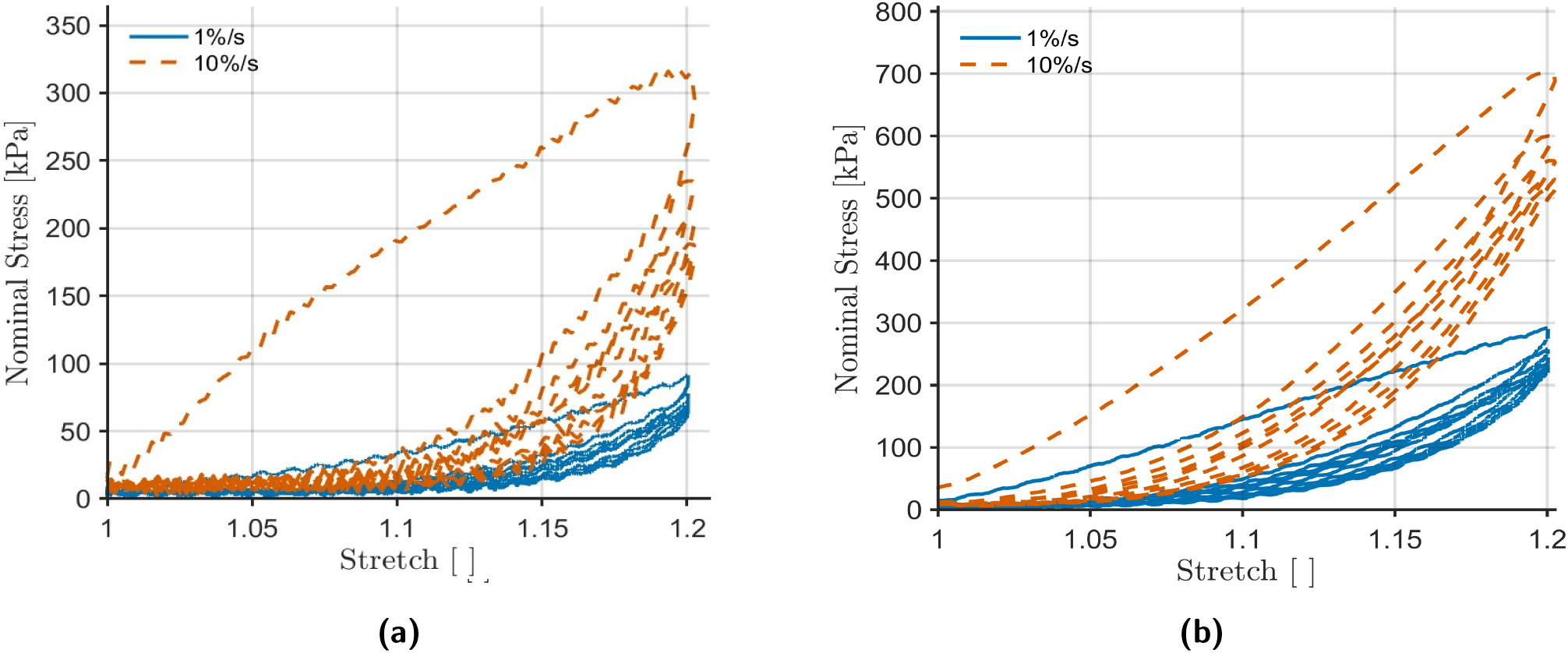
Representative stress–stretch responses of the masseter muscle samples under constant stretch-level cyclic loading at two strain rates (0.01 s^−1^ and 0.1 s^−1^). (a) Fresh samples and (b) formalin-preserved samples.

**Fig. 13:**
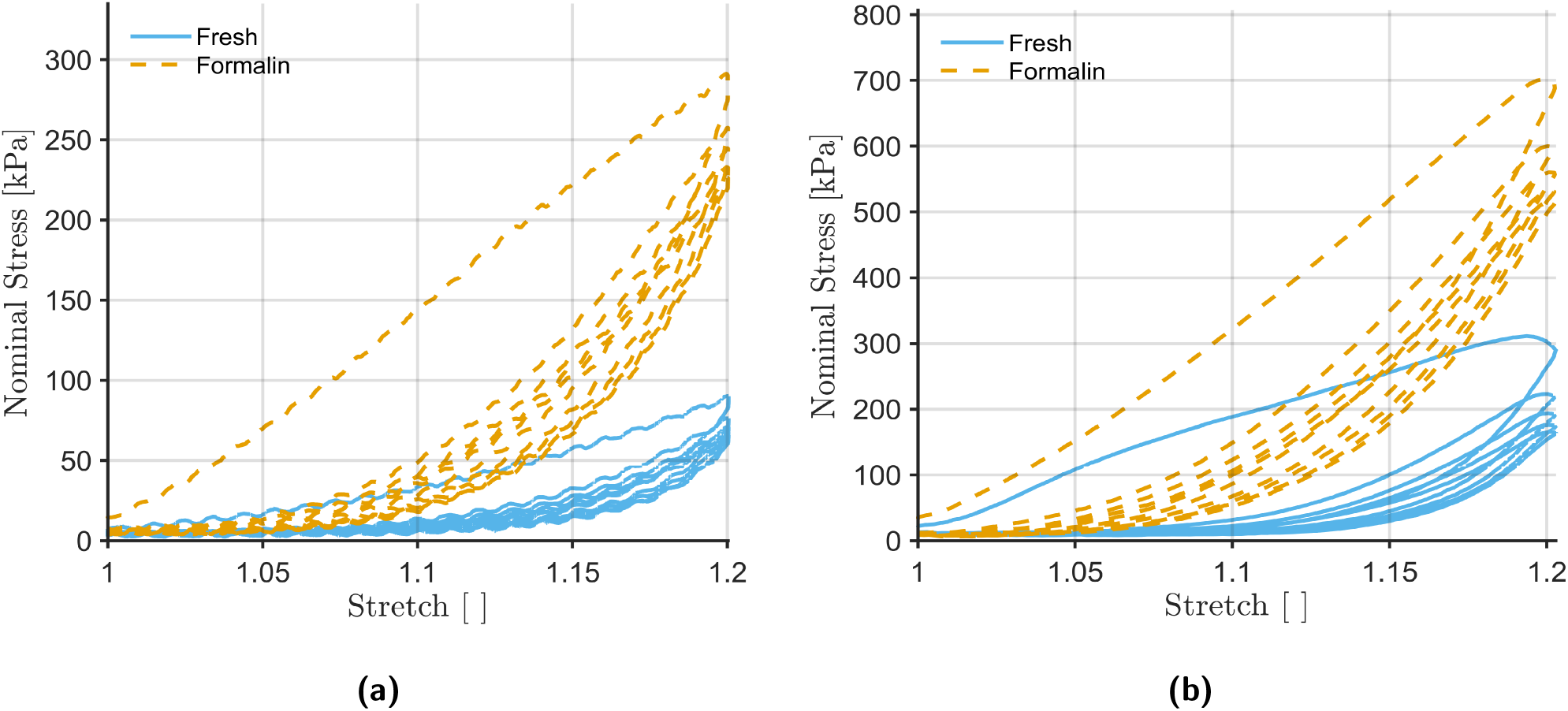
Representative stress–stretch responses of fresh and formalin-preserved masseter muscle samples under constant stretch-level cyclic loading at (a) 0.01 s^−1^ and (b) 0.1 s^−1^

#### 3.3.1. Strain rate-dependent behaviour

The response curves of fresh and formalin-preserved samples under continuous stretch protocol at different strain rates are shown in Fig. 10(a) and Fig. 10(b), respectively. The stress values are higher at a strain rate of 0.1 s^−1^ than at 0.01 s^−1^ across the entire stretch range. In the higher stretch region, the slope of the stress–stretch loading curve at 0.1 s^−1^ is steeper than that at 0.01 s^−1^. A similar trend is observed under the constant stretch protocol (Fig. 12(a) and Fig. 12(b)), where the stress levels during cyclic loading at 0.1 s^−1^ are consistently higher than those at 0.01 s^−1^. Therefore, for both continuous and constant stretch protocols, a rate-dependent effect is observed across all cycles.

#### 3.3.2. Fresh vs. formalin-preserved tissue behaviour

The stress–stretch responses of fresh and formalin-preserved samples under continuous stretch protocol at a strain rate of 0.01 s^−1^ are shown in Fig. 11(a), while those at 0.1 s^−1^ are shown in Fig. 11(b). From these figures, it can be observed that the formalin-preserved samples exhibit higher stress values than the fresh samples at corresponding stretch levels for both strain rates. In addition, the slope of the stress–stretch loading curves at 0.1 s^−1^ is steeper than that at 0.01 s^−1^ for both preservation conditions. A similar trend is observed for fresh and formalin-preserved tissues under constant stretch protocol (Fig. 13(a) and Fig. 13(b)). Among the four test conditions (fresh at 0.01 s^−1^, formalin-preserved at 0.01 s^−1^, fresh at 0.1 s^−1^, and formalin-preserved at 0.1 s^−1^), the highest stress values are observed for the formalin-preserved samples at 0.1 s^−1^, irrespective of the loading protocol.

### 3.4. Tangent modulus analysis

The tangent modulus is a measure of the stiffness of the tissue. For each sample, the tangent modulus was calculated from the first loading curve of the stress–stretch response between the stretch levels of 1.00 and 1.05. Fig. 14(a) and Fig. 14(b) show the boxplots illustrating the median, interquartile range (IQR), and variability of the tangent modulus for the four test conditions under continuous and constant stretch protocols, respectively. From Fig. 14, it can be observed that for both the continuous and constant stretch protocols, formalin-preserved samples exhibit higher median tangent modulus values than fresh samples at both strain rates. In addition, samples tested at a strain rate of 0.1 s^−1^ show higher tangent modulus values than those tested at 0.01 s^−1^ within each preservation condition. A larger spread in tangent modulus values for formalin-preserved samples, particularly at 0.1 s^−1^ can also be observed.

**Fig. 14:**
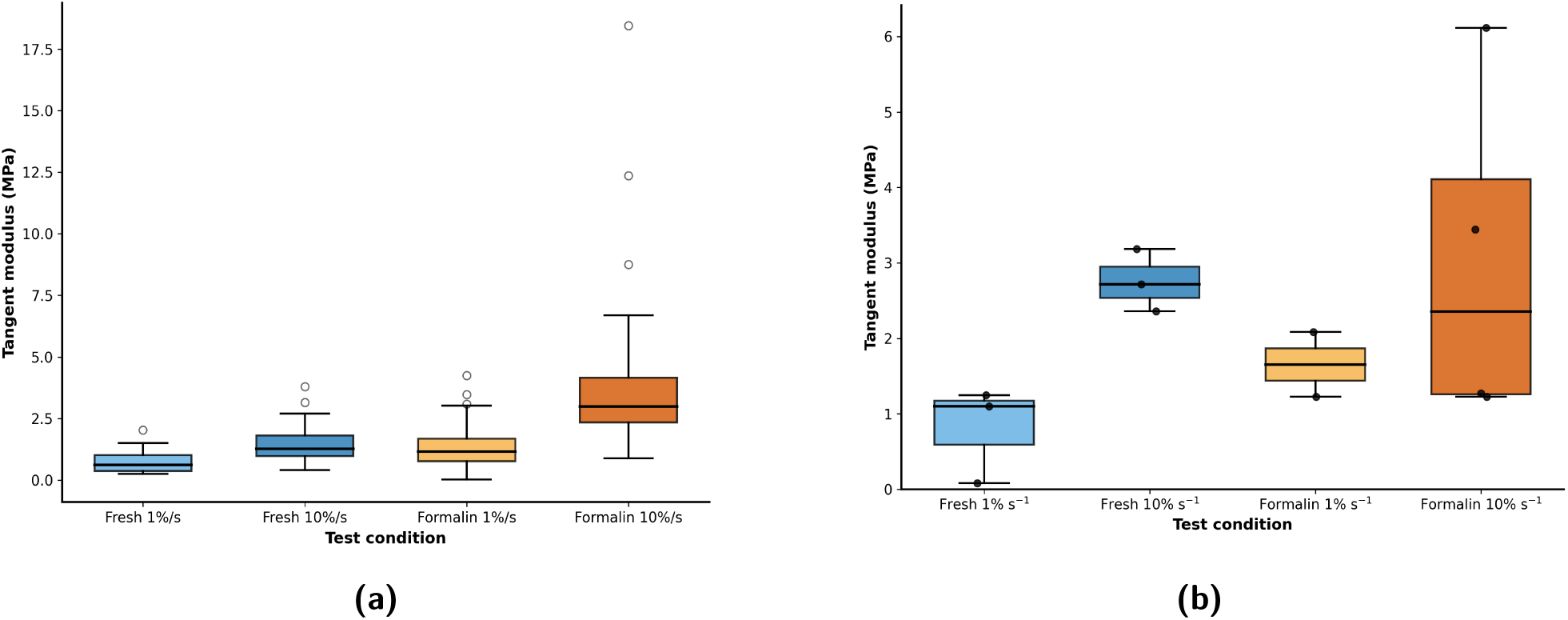
Boxplots of tangent modulus for the four test conditions (Fresh 0.01 s^−1^, Fresh 0.1 s^−1^, Formalin 0.01 s^−1^, Formalin 0.1 s^−1^) under the continuous stretch protocol (a) and the constant stretch protocol (b). The median (central line), interquartile range (box), and variability (whiskers and outliers) are shown.

Under the constant stretch protocol, the formalin-preserved samples at 0.1 s^−1^ exhibit a wide distribution of values, and the median does not show a clear increase compared to the corresponding fresh samples. This increased variability limits definitive comparison between groups.

### 3.5. Normalized hysteresis area (energy dissipation)

When the masseter muscle samples were subjected to cyclic uniaxial tensile testing, the obtained stress–stretch responses during loading–unloading cycles formed hysteresis loops, as shown in Fig. 12—Fig. 15. The energy dissipated during each cycle was quantified using the normalized hysteresis area, as described in Section 2.4.

**Fig. 15:**
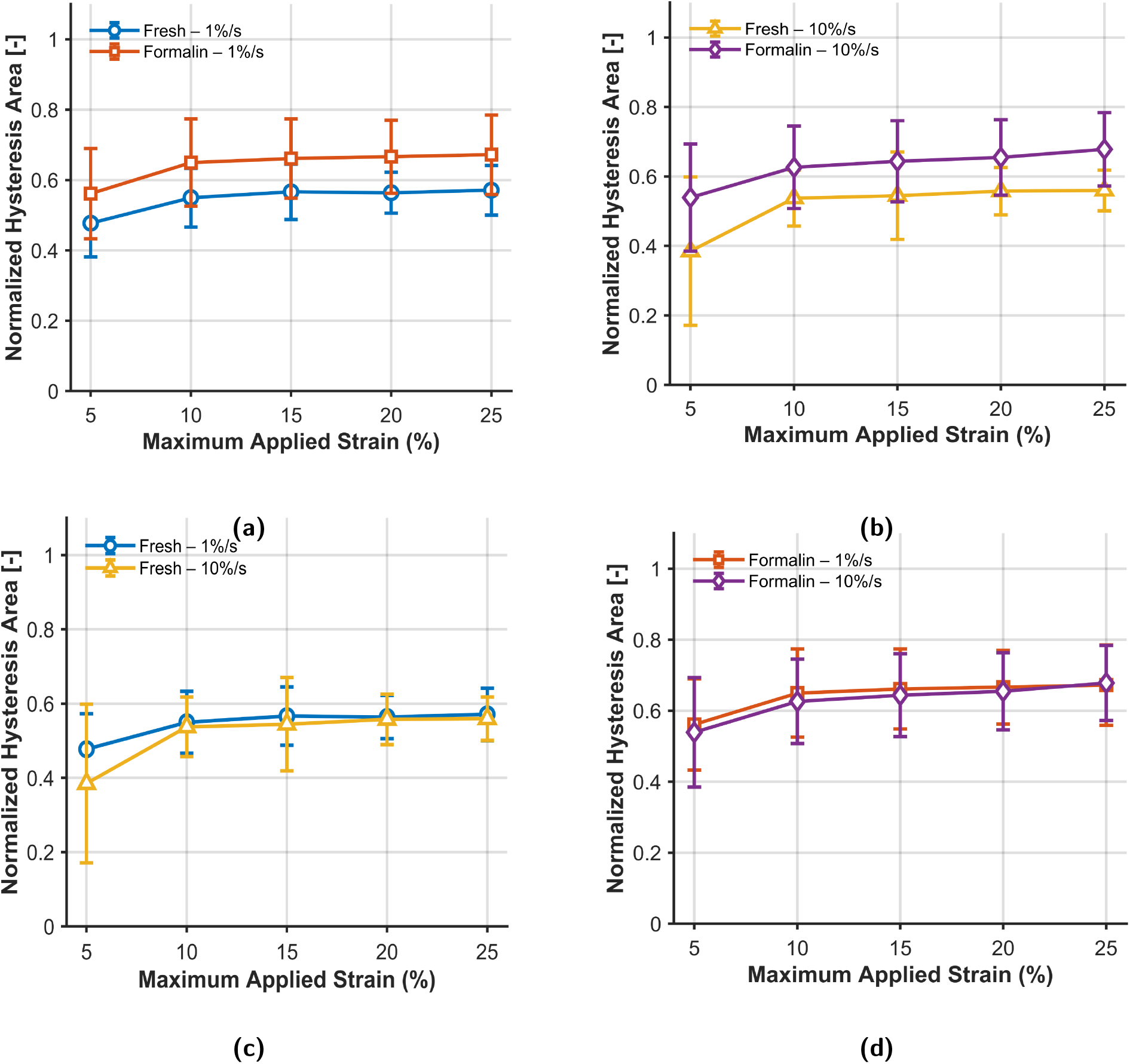
Comparison of normalized hysteresis area under the continuous stretch protocol at two strain rates. (a) Preservation effect at 0.01 s^−1^: fresh versus formalin-preserved samples. (b) Preservation effect at 0.1 s^−1^: fresh versus formalin-preserved samples. (c) Strain-rate effect in fresh samples: 0.01 s^−1^ versus 0.1 s^−1^. (d) Strain-rate effect in formalin-preserved samples: 0.01 s^−1^ versus 0.1 s^−1^. Data are presented as mean with standard deviation.

The variation in normalized hysteresis area with increasing maximum applied strain for the four test conditions under continuous stretch protocol is shown in Fig. 15 (a–d) and, under the constant stretch protocol is shown in Fig. 16(a-d). From Fig. 15 (a–d) , it can be observed that there is a steep increase in hysteresis values (normalized hysteresis area) during the initial applied strain between 5% and 10%. After that, the hysteresis values gradually increase with increasing applied strain and finally stabilize between 20% and 25% applied strain. A decrease in hysteresis values from the 1st cycle to 2nd cycle is observed for the constant stretch protocol Fig. 16(a-d), followed by a gradual decrease from the 2nd cycle to the 5th cycle. Under the continuous stretch protocol, at both strain rates (0.01 s^−1^, 0.1 s^−1^) formalin-preserved samples exhibited consistently higher hysteresis values than fresh samples (Fig. 15 (a) and (b)), while fresh samples exhibited higher hysteresis values than formalin-preserved samples under the constant stretch protocol (Fig. 16 (a) and (b)). For both preservation conditions, hysteresis values at a strain rate of 0.01 s^−1^ is slightly higher than those at 0.1 s^−1^ . (Fig. 15 (c) and (d) under the continuous stretch protocol, whereas under the constant stretch protocol, the hysteresis values at 0.1 s^−1^ are observed higher than those at 0.01 s^−1^ strain rate (Fig. 16 (c) and (d)). Overall, the normalized hysteresis area under continuous stretch protocol shows stretch-dependent pattern in all groups, characterized by an initial increase followed by stabilization at higher applied strain levels. On comparing the four test conditions under continuous stretch protocol, formalin-preserved samples exhibited greater normalized hysteresis area than fresh samples at the corresponding strain rates. Similarly, samples tested at 0.01 s^−1^ exhibited higher hysteresis values than those tested at 0.1 s^−1^ for both preservation conditions. Under the constant stretch protocol, the hysteresis area during the initial cycle was large, followed by a decrease in values in subsequent cycles, and reached stabilization at the final cycle.

**Fig. 16:**
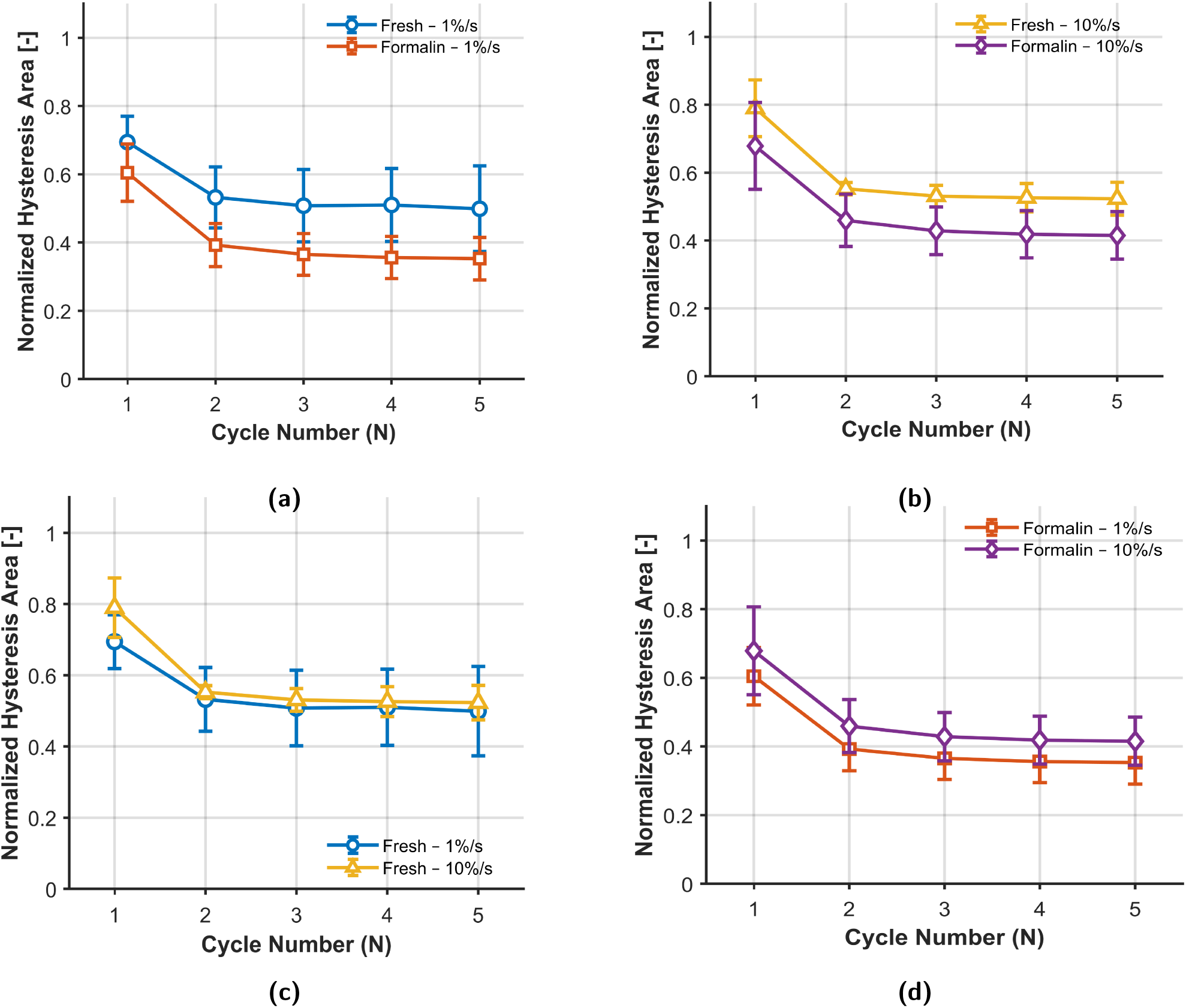
Comparison of normalized hysteresis area under the constant stretch protocol at two strain rates. (a) Preservation effect at 0.01 s^−1^: fresh versus formalin-preserved samples. (b) Preservation effect at 0.1 s^−1^: fresh versus formalin-preserved samples. (c) Strain-rate effect in fresh samples: 0.01 s^−1^ versus 0.1 s^−1^. (d) Strain-rate effect in formalin-preserved samples: 0.01 s^−1^ versus s^−1^. Data are presented as mean with standard deviation.

### 3.6. Stress-softening and cyclic conditioning behaviour

When the masseter muscle samples were subjected to cyclic uniaxial tensile testing with incrementally applied strain from 5% to 25% under the continuous stretch protocol, a reduction in stress at a given stretch level was observed during repeated loading, referred to as the Mullins effect or stress-softening, as shown in Fig. 10–Fig. 11. Under the constant stretch protocol, during repeated loading–unloading at a maximum applied strain of 20% for 5 cycles, a reduction in stress was observed in the subsequent cycles, and the tissue response eventually stabilized during the fourth and fifth cycles, referred to as cyclic conditioning (Fig. 12– Fig. 13). The stress-softening behaviour of the sample is quantified using softening index, which is measured as the variation in the strain energy density of the reloading curve with respect to the virgin curve (Equation.(9)). Fig. 17(a–d) shows the variation in the softening index for the four test conditions under the continuous stretch and Fig. 18(a–d) shows the variation in the conditioning index for the four test conditions under constant stretch protocol. Under the continuous stretch protocol, the softening index increases with increasing applied strain, indicating that the strain energy density difference increases with incremental applied strain (Fig. 17(a–d)). This reflects a progressive reduction in stress values during successive loading at higher stretch levels. For the four test conditions, during the initial increment in applied strain from 5% to 10%, a steep increase in the softening index is observed, followed by a gradual increase at higher applied strain levels. On examining the influence of preservation on softening behaviour, it can be observed that formalin-preserved samples exhibited higher softening behaviour than fresh samples for both strain rates (0.01 s^−1^, 0.1 s^−1^) (Fig. 17(a–b)). Similarly, when examining the effect of strain rate, it can be observed that the softening behaviour of the samples at a strain rate of 0.01 s^−1^ is higher than that at 0.1 s^−1^ (Fig. 17(c–d)), however, this effect is secondary to the influence of the maximum applied strain. A larger difference in softening behaviour between fresh and formalin-preserved samples can be observed at a strain rate of 0.1 s^−1^. Under the constant stretch protocol, the cyclic conditioning behaviour is quantified using the conditioning index, defined as the variation in the strain energy density of the reloading curve with respect to the first loading curve (Fig. 18(a–d)). A progressive increase in the conditioning index during successive loading cycles at the maximum applied strain is observed, as shown in Fig. 18(a–d).

**Fig. 17:**
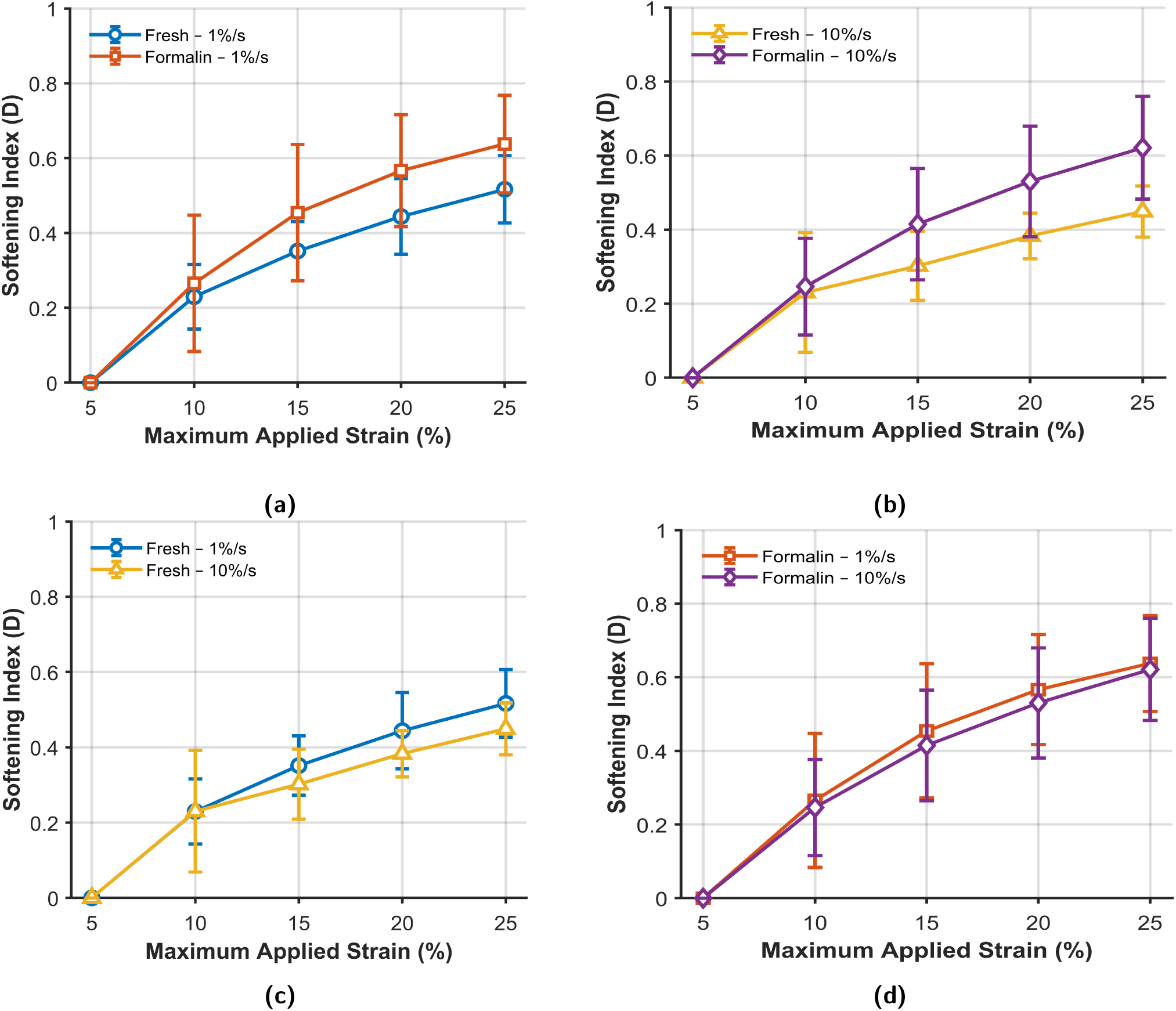
Comparison of stress-softening index under the continuous stretch protocol at two strain rates. (a) Preservation effect at 0.01 s^−1^: fresh versus formalin-preserved samples. (b) Preservation effect at 0.1 s^−1^: fresh versus formalin-preserved samples. (c) Strain-rate effect in fresh samples: 0.01 s^−1^ versus 0.1 s^−1^. (d) Strain-rate effect in formalin-preserved samples: 0.01 s^−1^ versus s^−1^. Data are presented as mean with standard deviation.

**Fig. 18:**
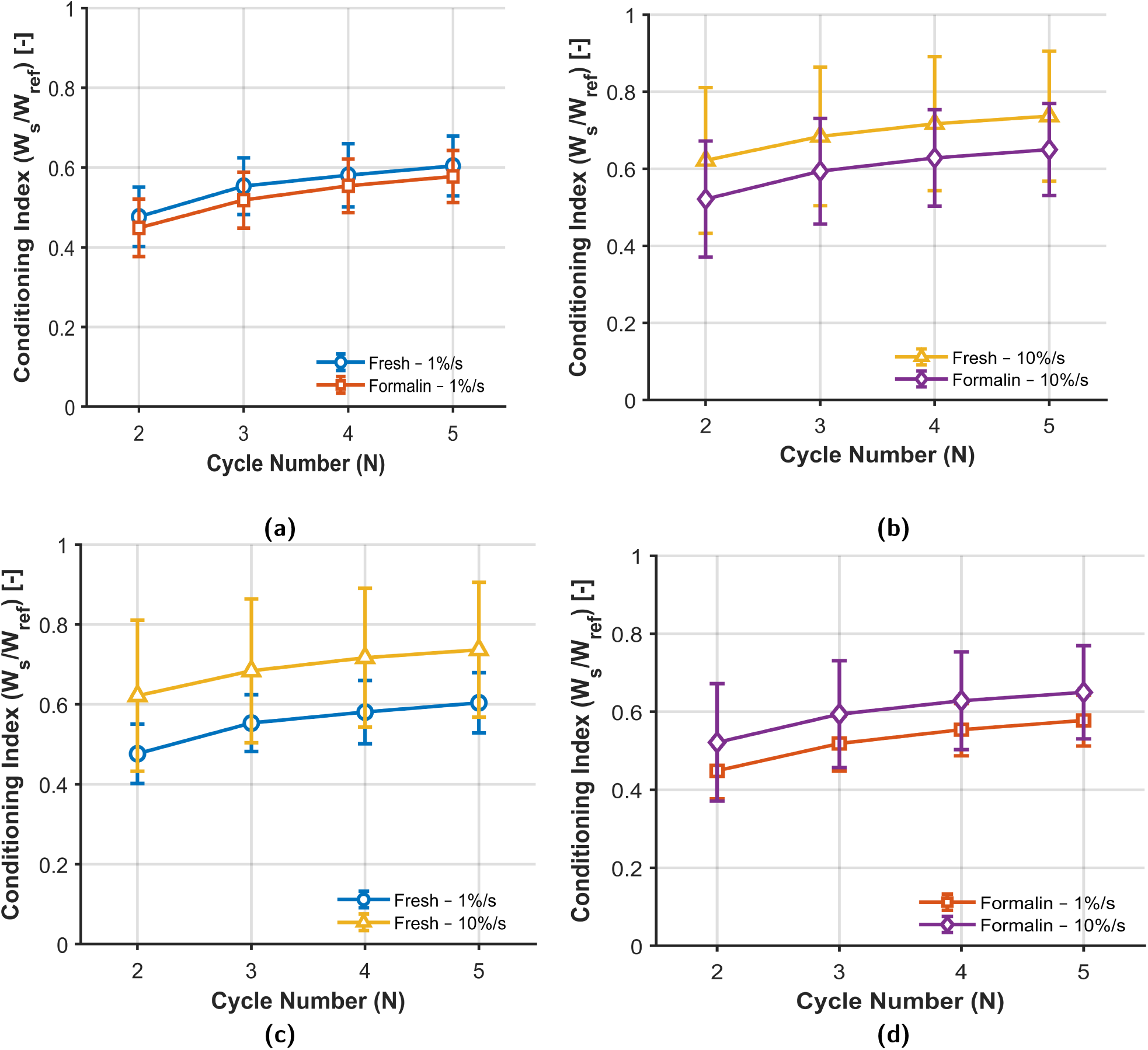
Comparison of conditioning under the constant stretch protocol at two strain rates. (a) Preservation effect at 0.01 s^−1^: fresh versus formalin-preserved samples. (b) Preservation effect at 0.1 s^−1^: fresh versus formalin-preserved samples. (c) Strain-rate effect in fresh samples: 0.01 s^−1^ versus 0.1 s^−1^. (d) Strain-rate effect in formalin-preserved samples: 0.01 s^−1^ versus 0.1 s^−1^. Data are presented as mean with standard deviation.

### 3.7. Residual strain

When the masseter muscle samples were subjected to cyclic uniaxial tensile testing with incrementally applied train levels from 5% to 25% under the continuous stretch protocol, permanent tissue deformation was observed during unloading. This is characterized by a non-zero strain at which the stress returns to zero, referred to as the residual strain (Fig. 10–Fig. 13). The residual strain was calculated at each stretch level from the unloading curve. Under the constant stretch protocol, during repeated loading–unloading at a maximum applied strain of 20% for 5 cycles, residual strain values were observed during unloading in each cycle (Fig. 14–Fig. 15). Under the continuous stretch protocol, the residual strain increases with increasing applied strain, indicating the accumulation of permanent deformation (Fig. 19(a–d)). For the four test conditions, gradual increase in the residual strain was observed. On examining the influence of preservation on softening behaviour, it can be observed that formalin-preserved samples exhibited higher residual strain values than fresh samples for both strain rates (0.01 s^−1^, 0.1 s^−1^) (Fig. 19(a–b)). Similarly, when examining the effect of strain rate, it can be observed that the residual strain values at a strain rate of 0.01 s^−1^ are higher than that at 0.1 s^−1^ (Fig. 19(c–d)). A larger difference in residual strain values between fresh and formalin-preserved samples can be observed at a strain rate of 0.01 s^−1^. Under the constant stretch protocol, higher values of residual strain were observed during first and second cycles, after which there was a gradual increase in strain values for all the test conditions (Fig. 20(a–d)). At both the strain rates, fresh samples exhibited higher residual strain values than formalin-preserved samples (Fig. 20(a–b)).

**Fig. 19:**
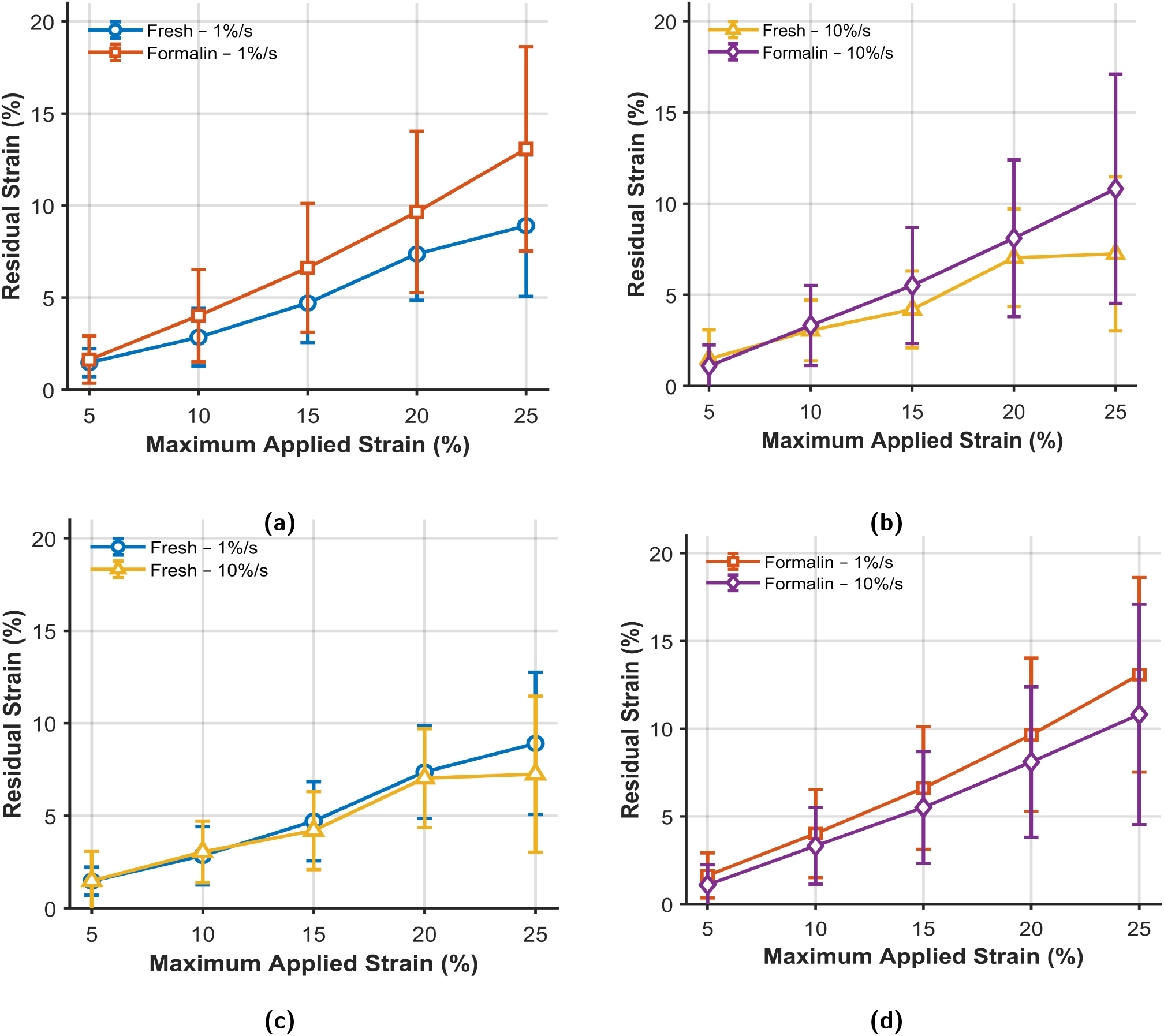
Comparison of residual strain under the continuous stretch protocol at two strain rates. (a) Preservation effect at 0.01 s^−1^: fresh versus formalin-preserved samples. (b) Preservation effect at 0.1 s^−1^: fresh versus formalin-preserved samples. (c) Strain-rate effect in fresh samples: 0.01 s^−1^ versus 0.1 s^−1^. (d) Strain-rate effect in formalin-preserved samples: 0.01 s^−1^ versus 0.1 s^−1^. Data are presented as mean with standard deviation.

**Fig. 20:**
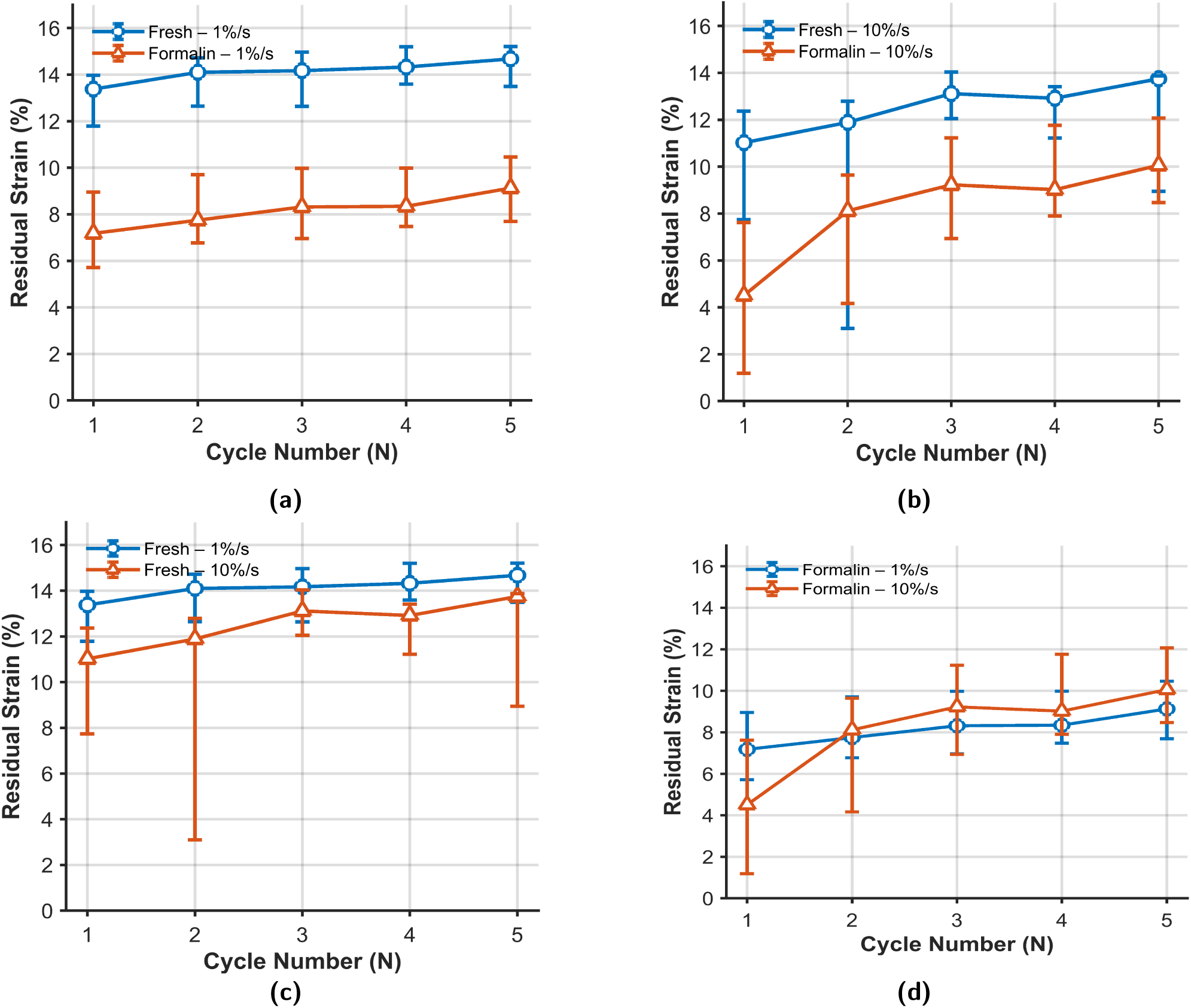
Comparison of residual strain under the constant stretch protocol at two strain rates. (a) Preservation effect at 0.01 s^−1^: fresh versus formalin-preserved samples. (b) Preservation effect at 0.1 s^−1^: fresh versus formalin-preserved samples. (c) Strain-rate effect in fresh samples: 0.01 s^−1^ versus 0.1 s^−1^. (d) Strain-rate effect in formalin-preserved samples: 0.01 s^−1^ versus 0.1 s^−1^. Data are presented as mean with standard deviation.

### 3.8. Statistical comparison of mechanical properties

The statistical pairwise comparisons between fresh and formalin-preserved samples at both strain rates, as well as between 0.01 s^−1^ and 0.1 s^−1^ within each preservation condition, were performed for the corresponding features, including tangent modulus, normalized hysteresis area, stress-softening (conditioning) index, and residual strain using the Mann–Whitney U test with a significance level of *p* < 0.05.

For the continuous stretch protocol, statistically significant differences (*p* < 0.05) were observed between fresh and formalin-preserved samples at both strain rates for all the considered features. In contrast, for the strain-rate comparisons within each preservation condition, statistically significant differences were observed only for the tangent modulus, whereas normalized hysteresis area, stress-softening index, and residual strain did not show statistically significant differences (*p* > 0.05).

Under the constant stretch protocol, no statistically significant differences (*p* > 0.05) were observed for any of the features. This may be due to the limited sample size in each group, which reduces the statistical power of the test.

The statistical results are presented in Table 6.

**Table 6.** Statistical comparison of mechanical characteristics between test conditions under continuous and constant stretch protocols using the Mann–Whitney U test. Statistically significant group (*p* < 0.05) and its corresponding *p*-values are presented for each characteristic. “–” indicates no statistically significant difference (*p* > 0.05).

| Comparison | Group 1 | Group 2 | Tangent Modulus | Normalized Hysteresis Area | Softening Index | Residual Strain |
| --- | --- | --- | --- | --- | --- | --- |
| **Continuous Stretch Protocol** |  |  |  |  |  |  |
| Fresh vs Formalin | Fresh 0.01 s<sup>-1</sup> | Formalin 0.01 s<sup>-1</sup> | Formalin 0.01 s<sup>-1</sup><br>7.82 × 10<sup>-4</sup> | Formalin 0.01 s<sup>-1</sup><br>1.39 × 10<sup>-4</sup> | Formalin 0.01 s<sup>-1</sup><br>3.54 × 10<sup>-4</sup> | Formalin 0.01 s<sup>-1</sup><br>9.85 × 10<sup>-4</sup> |
|  | Fresh 0.1 s<sup>-1</sup> | Formalin 0.1 s<sup>-1</sup> | Formalin 0.1 s<sup>-1</sup><br>1.96 × 10<sup>-5</sup> | Formalin 0.1 s<sup>-1</sup><br>2.64 × 10<sup>-5</sup> | Formalin 0.1 s<sup>-1</sup><br>2.63 × 10<sup>-4</sup> | Formalin 0.1 s<sup>-1</sup><br>0.0274 |
| 0.01 s<sup>-1</sup> vs 0.1 s<sup>-1</sup> | Fresh 0.01 s<sup>-1</sup> | Fresh 0.1 s<sup>-1</sup> | Fresh 0.1 s<sup>-1</sup><br>5.86 × 10<sup>-4</sup> | – | – | – |
|  | Formalin 0.01 s<sup>-1</sup> | Formalin 0.1 s<sup>-1</sup> | Formalin 0.1 s<sup>-1</sup><br>8.33 × 10<sup>-8</sup> | – | – | – |
| **Constant Stretch Protocol** |  |  |  |  |  |  |
| Fresh vs Formalin | Fresh 0.01 s<sup>-1</sup> | Formalin 0.01 s<sup>-1</sup> | – | – | – | – |
|  | Fresh 0.1 s<sup>-1</sup> | Formalin 0.1 s<sup>-1</sup> | – | – | – | – |
| 0.01 s<sup>-1</sup> vs 0.1 s<sup>-1</sup> | Fresh 0.01 s<sup>-1</sup> | Fresh 0.1 s<sup>-1</sup> | – | – | – | – |
|  | Formalin 0.01 s<sup>-1</sup> | Formalin 0.1 s<sup>-1</sup> | – | – | – | – |

## 4. Discussion

The experimental results characterized the human masseter muscle as nonlinear and viscoelastic material. The experimental findings from this study are compared to provide new insight into the nonlinear, time-dependent, and history-dependent mechanical behaviour of the human masseter muscle under cyclic uniaxial tensile testing, particularly with respect to the effects of strain rate, tissue preservation condition, and testing protocol. Furthermore, the experimental results obtained are related to the histological findings, physiological functions and the previous experimental studies on masseter muscle. The representative stress-stretch response of the tissue under continuous and constant protocol (Fig. 10–Fig. 13) exhibit a J-shaped response in the loading branch, which is consistent with the histological study of the human masseter muscle. The initial non-linear response, followed by a quasi-linear region and a final nonlinear stiffening at higher stretch levels are due to the heterogenous fibre distribution of slow-twitch (Type I), fast-twitch (Type II), and intermediate fibres [41]. Overall, the masseter muscle becomes progressively stiffer with increasing stretch, thereby exhibiting nonlinear behaviour. During the initial loading phase at lower stretch levels, the crimped collagen fibers begin to straighten, which results in a small rise in stress values (toe region). On increasing the stretch levels further, the collagen fibers become fully aligned and the load is transferred along the fiber direction, resulting in quasilinear increase in stress (intermediate region). At higher stretch levels, the fibres are fully aligned and stretched, resulting in a nonlinear stiffening response, where the tissue exhibits a steeper increase in stress [42]. This nonlinear behaviour is also consistent with the physiological function of the masseter muscle, which is subjected to combined mechanical solicitation and compression during mastication. This finding is in agreement with the ex vivo uniaxial tensile study conducted by Picard et al. [31], who reported a similar nonlinear stress–strain response for facial soft tissues, including the masseter muscle. Furthermore, the present results are in agreement with previous experimental studies on the nonlinear mechanical behaviour of facial soft tissues [11, 12, 16, 18].

A large variation in the stress–stretch response and failure response (Fig. 8) of the samples was observed. It should be noted that, according to the different loading protocols employed in this study, the responses are not purely monotonic and therefore do not correspond directly to true rupture behaviour. The observed variability is attributed to the heterogeneous and complex fibre structure of the masseter muscle, which is consistent with anatomical and histological findings [43, 44]. The inter-sample variability also arises from donor demographic factors such as age and gender, as supported by recent studies investigating the influence of age [45] and gender [29] on the mechanical properties of facial soft tissues. The use of a distribution-based statistical approach (Shapiro–Wilk test followed by Fréchet fitting) to select representative curves is therefore justified. The right-skewed distribution of tangent modulus (Fig.14) confirms that the mechanical properties of the masseter muscle do not follow a normal distribution, which is consistent with findings in other soft tissues [46, 36].

### Strain rate-dependent behaviour

From the experimental results, a clear strain-rate-dependent response of the human masseter muscle under both continuous and constant stretch protocols (Fig. 10–Fig. 12) can be observed. The stiffness increases at a higher strain rate (0.1 % s^−1^) when compared to 0.01 % s^−1^. In all regions of the stress–stretch curves, the slope at 0.1 % s^−1^ is higher than at 0.01 % s^−1^. This behaviour is further supported quantitatively by the tangent modulus, which increases with strain rate (Fig. 14). This time-dependent response confirms the viscoelastic nature of the masseter muscle. The results are consistent with previous experimental studies on soft biological tissues. Har-Shai et al. [47] studied the viscoelastic behaviour of the SMAS under varying loading rates and reported that soft tissue deformation is strongly influenced by loading rate and duration. Chawla et al. [48] showed that human soft tissues are strain-rate dependent, and that increasing the strain rate alters the stress–strain response. Soft tissues exhibit increased stiffness at higher strain levels with increasing strain rate. From the microstructural perspective, the increase in stiffness at higher strain rates may be due to the progressive alignment of the collagen fibres [41] and the increase in collagen cross-link density. Our findings were also supported by the study conducted by [36], who reported that the density of collagen cross-link has a significant influence on the tissue stiffness. Also, the complex fibre architecture of the masseter muscle [41] contributes to the variation in mechanical response of the tissue under different strain rates.

The present study also shows the dependence of the hysteresis behaviour of the masseter muscle tissue on the applied strain and the strain rate. The normalized hysteresis, increases with increasing stretch levels, while it decreases with increased strain rate under the continuous stretch protocol. Under repeated loading, the tissue undergoes microstructural rearrangements and stabilizes, exhibiting stable energy dissipation, referred to as cyclic conditioning. The hysteresis behaviour observed in this study is consistent with the previously reported viscoelastic responses of biological soft tissues [49]. Durcan et al. [36] performed continuous stretch cyclic tests and quantified hysteresis for each loading–unloading cycle, with the results indicating a dependence of hysteresis behaviour on the applied stretch level and strain rate. Fung [50] described that the stress response of soft biological tissues not only depends on the applied strain but also on the rate of loading, indicating time-dependent behaviour. Under the constant stretch protocol, the hysteresis decreases with increasing cycle number under repeated loading, while it increases with increasing strain rate. A similar behaviour has been observed in previous studies on biological soft tissues. Remache et al. [38] showed that, during cyclic tensile loading of porcine skin, the hysteresis area gradually decreases with repeated loading. Har-Shai et al. [47] similarly reported reduced hysteresis with repeated cyclic loading in SMAS tissue.

The strain-rate dependence on stress-softening was observed under continuous stretch protocol. The softening index at 0.01 % s^−1^ is higher than at 0.1 % s^−1^ for both fresh and formalin-preserved tissues. Under constant stretch protocol, the conditioning index at 0.1 % s^−1^ is higher than at 0.01 % s^−1^ for both fresh and formalin-preserved tissues. A similar interpretation on strain rate-dependent behaviour is presented by Durcan et al. [36],Anssari-Benam et al. [51],Haldar and Pal [52] and Fung [50], who reported that the stress response of soft tissues depends on loading rate and applied strain.

The effect of strain rate on residual strain behaviour is evident in this study. Under the continuous stretch protocol, the residual strain increases gradually with increasing applied strain for all four test conditions. This behaviour is due to the complex fiber architecture of the masseter muscle tissue. With increasing stretch levels, progressive fibre recruitment occurs along with the accumulation of structural damage, resulting in a non-zero strain after unloading. Thus, the increase in residual strain with increasing stretch levels is consistent with the history-dependent behaviour of the masseter muscle. The residual strain at 0.01 % s^−1^ is higher than at 0.1 % s^−1^ and the results are consistent with previous studies on biological soft tissues reported by Durcan et al. and Fung [50] on the history-dependent mechanical behaviour of soft tissues.

### Effects of preservation (fresh vs formalin)

The formalin-preserved samples exhibit higher stiffness than fresh samples at both strain rates and under both loading conditions (Fig. 11, Fig. 13). The tangent modulus analysis also shows that formalin-preserved samples exhibit higher median values at both strain rates (Fig. 14).

This behaviour is supported by previous studies, which reported increased stiffness in embalmed soft tissues. For instance, Joy et al. [30] reported stiffness values (Young’s modulus) for various fresh and embalmed soft tissues, including the masseter muscle, using in vivo shear wave elastography. Their investigations show that Thiel-embalmed masseter tissue exhibits higher stiffness than that of human volunteers. The ex vivo study on fresh and preserved tendons by Hohmann et al. [34] reported that formalin preservation has a significant influence on the mechanical properties of the tissue. The difference in fresh and formalin-preserved tissue behaviour is also attributed to its microstructural organization, where the increased stiffness may be due to an increase in collagen cross-linking caused by formalin. This is also supported by the studies conducted by Durcan et al. [36] and Hohmann et al. [34] who reported that embalming increases the amount of cross-linking of collagen fibres, thereby altering the magnitude of the mechanical properties of soft tissues.

The effect of preservation on hysteresis behaviour is also evident in this study and is consistent with previous studies on the influence of preservation on the mechanical behaviour of soft tissues. As stated, formalin increases the collagen cross-link density, thereby influencing the viscoelastic behaviour of the tissue. Under continuous stretch protocol, the formalin-preserved samples exhibit higher stress-softening at both strain rates. The results are consistent with the study conducted by Durcan et al. [36] who compared fresh and formalin-preserved oesophageal tissue and presented that formal-pre served tissues exhibit higher magnitude of stress-softening due to the increase in collagen cross-link density.

Sample preservation influences the cyclic conditioning of the tissue, with formalin-preserved samples exhibiting a lower conditioning index than fresh samples. Due to repeated loading to a constant stretch level, the increase in collage cross-link density of formalin-preserved samples limits internal structural rearrangements leading to lower conditioning index than fresh samples. A similar study on the effect of preservation on stress-softening was conducted by Caro-Bretelle et al. [53] preservation modifies the collagen network and alters the stress-softening and damage behaviour of soft tissues

The effect of preservation on residual strain behaviour is evident in this study. Under the continuous stretch protocol, the formalin-preserved samples exhibit higher residual strain than the fresh samples, as formalin increases the collagen cross-link density. Similarly, the residual strain at 0.01 % s^−1^ is higher than at 0.1 % s^−1^ due to the greater time available for internal structural rearrangement and fluid redistribution. The results are consistent with previous studies on biological soft tissues reported by Durcan et al. [36] and Fung [50] on the history-dependent mechanical behaviour of soft tissues. Under the constant stretch protocol, repeated loading to a fixed maximum stretch also produces permanent deformation, observed as residual strain in every cycle. The residual strain of fresh samples is higher than that of formalin-preserved samples at both strain rates. This is supported in the study conducted by Caro-Bretelle et al. [53], who reported that preservation modifies permanent set through microstructural alteration.

The Mann–Whitney U test results (Table 6) revealed that, under the continuous stretch protocol, statistically significant differences (*p* < 0.05) were observed between fresh and formalin-preserved samples for all the considered characteristics, including tangent modulus, normalized hysteresis area, stress-softening index, and residual strain. This indicates that formalin preservation has a pronounced effect not only on the stiffness but also on the viscoelastic and history-dependent mechanical behaviour of the tissue.

In contrast, the strain-rate-dependent comparisons within each preservation condition showed statistically significant differences only for the tangent modulus, whereas normalized hysteresis area, stress-softening index, and residual strain did not exhibit statistically significant differences (*p* > 0.05). This suggests that strain rate predominantly influences the instantaneous stiffness of the tissue, while its effect on energy dissipation, stress-softening, and permanent deformation is less significant. Under the constant stretch protocol, no statistically significant differences (*p* > 0.05) were observed between the groups for all the characteristics. This might be due to the limited sample size used for this protocol, which may not be sufficient to determine differences in the characteristics between the groups.

## 5. Conclusion

In this ex vivo study, the nonlinear, and viscoelastic behaviour of the human masseter muscle was characterized using uniaxial cyclic tensile tests. The results of this study contribute to a better understanding of the relationship between the tissue’s mechanical properties, anatomical structure, and physiological function. Overall, the stress-stretch response of the masseter muscle exhibits nonlinear and viscoelastic behaviour. Additionally, the history-dependent behaviour of the tissue including stress-softening and residual strain, was also investigated. Under both continuous and constant stretch loading protocols, the mechanical behaviour of the masseter muscle including stiffness, hysteresis, stress-softening and residual strain, was found to be strongly influenced by applied strain, strain rate and formalin preservation, due to the changes in the internal microstructure of the tissue, caused by these factors.

Overall, this work provides new insights from the experimental findings on the non linear and viscoelastic behaviour of human masseter muscle, which has not been sufficiently characterized in previous ex vivo studies. The findings provide useful mechanical data for biomechanical modelling of the human masticatory system. Future work will focus on the development and validation of constitutive models for the human masseter muscle based on the experimental data presented in this study, with applications in cranio-maxillofacial surgical simulation, prosthetic design, and facial soft-tissue biomechanics.

